# A positive association between population genetic differentiation and speciation rates in New World birds

**DOI:** 10.1101/085134

**Authors:** Michael G. Harvey, Glenn F. Seeholzer, Brian Tilston Smith, Daniel L. Rabosky, Andrés M. Cuervo, John T. Klicka, Robb T. Brumfield

## Abstract

Although an implicit assumption of speciation biology is that population differentiation is an important stage of evolutionary diversification, its true significance remains largely untested. If population differentiation within a species is related to its speciation rate over evolutionary time, the causes of differentiation could also be driving dynamics of organismal diversity across time and space. Alternatively, geographic variants might be short-lived entities with rates of formation that are unlinked to speciation rates, in which case the causes of differentiation would have only ephemeral impacts. Combining population genetics datasets including 17,746 individuals from 176 New World bird species with speciation rates estimated from phylogenetic data, we show that the population differentiation rates within species predict their speciation rates over long timescales. Although relatively little variance in speciation rate is explained by population differentiation rate, the relationship between the two is robust to diverse strategies of sampling and analyzing both population-level and species-level datasets. Population differentiation occurs at least three to five times faster than speciation, suggesting that most populations are ephemeral. Population differentiation and speciation rates are more tightly linked in tropical species than temperate species, consistent with a history of more stable diversification dynamics through time in the Tropics. Overall, our results suggest investigations into the processes responsible for population differentiation can reveal factors that contribute to broad-scale patterns of diversity.

## Significance

The causes of differentiation among populations are well circumscribed, but it remains unclear if they impact the proliferation of organisms over deep time. If, as some recent theory and observations suggest, population differentiation is untethered from species formation, then the causes of population differentiation are unlikely to have long-term evolutionary effects. We provide the first large-scale test of the link between standardized estimates of rates of population differentiation from population genetic data and speciation rates. We find that relative population differentiation rates predict speciation rates across New World birds, confirming the potential macroevolutionary importance of causes of differentiation. We also find that population differentiation and speciation rates are more tightly linked in the Tropics, which may contribute to greater tropical species richness.

Speciation in most organisms is initiated via the geographic isolation and differentiation of populations. The rate that populations differentiate within a species is determined by many extrinsic and intrinsic factors, including the rate of geological and climatic change (1, 2), the dispersal ability of organisms (3), and the availability of ecological opportunities and strength of natural selection (4, 5). An implicit assumption of speciation biology is that these factors have an impact that percolates through to long evolutionary timescales and influences the proliferation of species. This connection, however, is not assured. The factors responsible for population differentiation can only affect species diversification if differentiation acts as a limiting control on diversification - for example, if the rate at which differentiated populations form within a given lineage determines the rate at which species can form in that lineage - or if differentiation and diversification are both responses to the same causal processes. In either case, differentiation and diversification should be associated across evolutionary lineages.

Recent work, however, suggests that differentiation dynamics are untethered from those of diversification. Some macroevolutionary biologists suggest that geographic populations or variants are often short-lived entities and that their rate of formation within a species might have little relation to speciation rates (6, 7, 8). Instead, speciation may be limited by other population-level processes, such as the persistence of differentiated populations (9) or the evolution of sufficient ecological divergence (5, 10) or reproductive isolation (11, 12) for differentiated populations to coexist in sympatry, or speciation may be random with respect to population-level processes. If differentiation is not associated with diversification, the factors responsible for differentiation cannot be expected to have macroevolutionary impacts.

The association between population differentiation and diversification can be tested by comparing present-day population differentiation within species to the speciation rate of their lineages over deeper evolutionary time. An association between differentiation and speciation rate across lineages would support the hypothesis that differentiation is important for speciation. Few studies have attempted to make this comparison (13, 14). Haskell and Adhikari (15) compared the number of taxonomic subspecies within species to the number of species in avian genera and found a correlation, suggesting that levels of geographic differentiation are a predictor of species diversity. Phillimore (16) used species stem ages from phylogenies to calculate the rate that their taxonomic subspecies had formed and found that the rate that subspecies accrued was correlated with phylogenetic speciation rates. Kisel and Barraclough (17), however, found no link between the magnitude of genetic divergence between populations and diversification rates in five sister clades of Costa Rican orchids. This conflicting evidence suggests that resolving the connection between differentiation and speciation is challenging, and will likely require the examination of standardized, quantitative metrics of population differentiation from a large sample of species.

Here, we assess the association between population differentiation and speciation rate using estimates of differentiation from population-level genetic data. We estimate population differentiation based on an application of the coalescent model to gene trees estimated from new and existing population genetic and phylogeographic datasets from 176 species of New World birds. We compare population differentiation to speciation rates estimated for the lineages subtending the same 176 species from phylogenetic trees of all birds. We first test whether population differentiation and speciation rates are associated across all sampled species. Because the association between population differentiation and speciation rates may vary across geographic contexts, we also test whether the association between differentiation and speciation rates in the Tropics differs from that in the Temperate Zone. Finally, we perform a suite of supplementary tests to assess the robustness of recovered relationships to the approach used for sampling and analysis.

## Results

We estimated population differentiation based on genetic data from range wide samples of individuals (*n* = 17,746) in 176 bird species from across the avian tree of life (Fig. 1*A*; *SI Materials and Methods*) and inhabiting all biogeographic regions of the New World (Fig. 1*B*). We used a Bayesian implementation of the Generalized Mixed Yule Coalescent model (18, 19) to standardize population differentiation estimates within each species (Fig. 1*C*). The number of genetically distinguishable geographic populations within species varied from one to 35 with a median of three (Fig. 2). Because species might vary in the number of geographic populations simply due to differences in age, we calculated the rate of population formation since the crown age of each species (age of the most recent common ancestor of extant haplotypes within the species) based on a time-calibrated phylogenetic tree. The rate at which geographic populations arose, hereafter the rate of population differentiation, varied from zero to 6.64 divergences/million years (My) with a median of 0.53 divergences/My (Fig. 2).

**Fig. 1.**
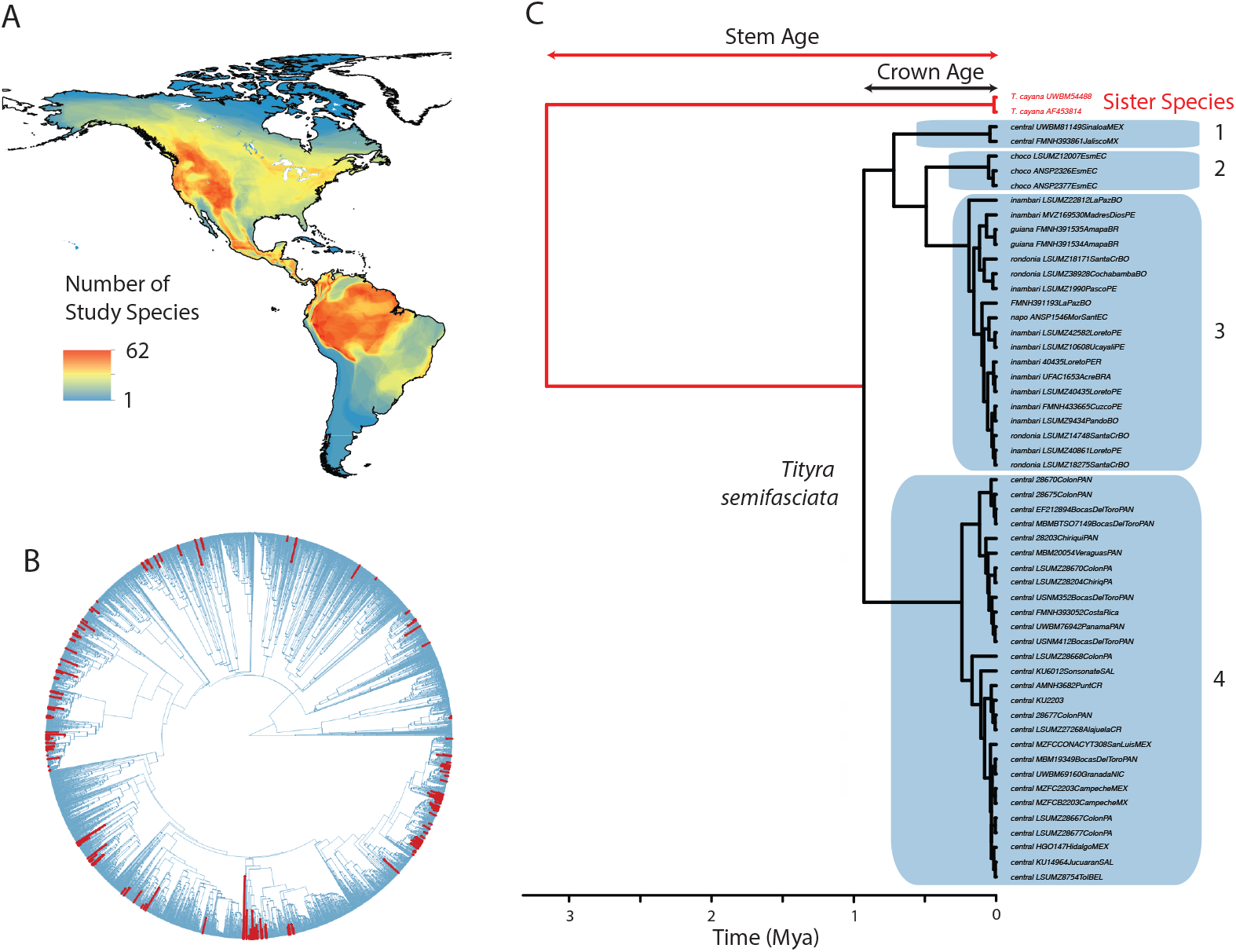
Sampling strategy and approach to measuring population differentiation. (A) Overlaid distribution maps from the New World bird species used to estimate amount of population genetic differentiation (*n* = 176). (B) The phylogenetic distribution of the study species within the tree of life of all birds (20). The red branches, which indicate the species examined in this study, occur throughout the tree and represent replicates with varying levels of phylogenetic independence for the purpose of comparative analysis. (C) An example of a mitochondrial gene tree used to estimate the rate of population genetic differentiation within one of the 176 study species, the Masked Tityra (*Tityra semifasciata*). The blue polygons represent population clusters for this species as inferred using bGMYC (19) based on a posterior probability threshold of shared population membership of 0.8. The stem age and crown age for this species, used to estimate rates of differentiation, are also depicted.

**Fig. 2.**
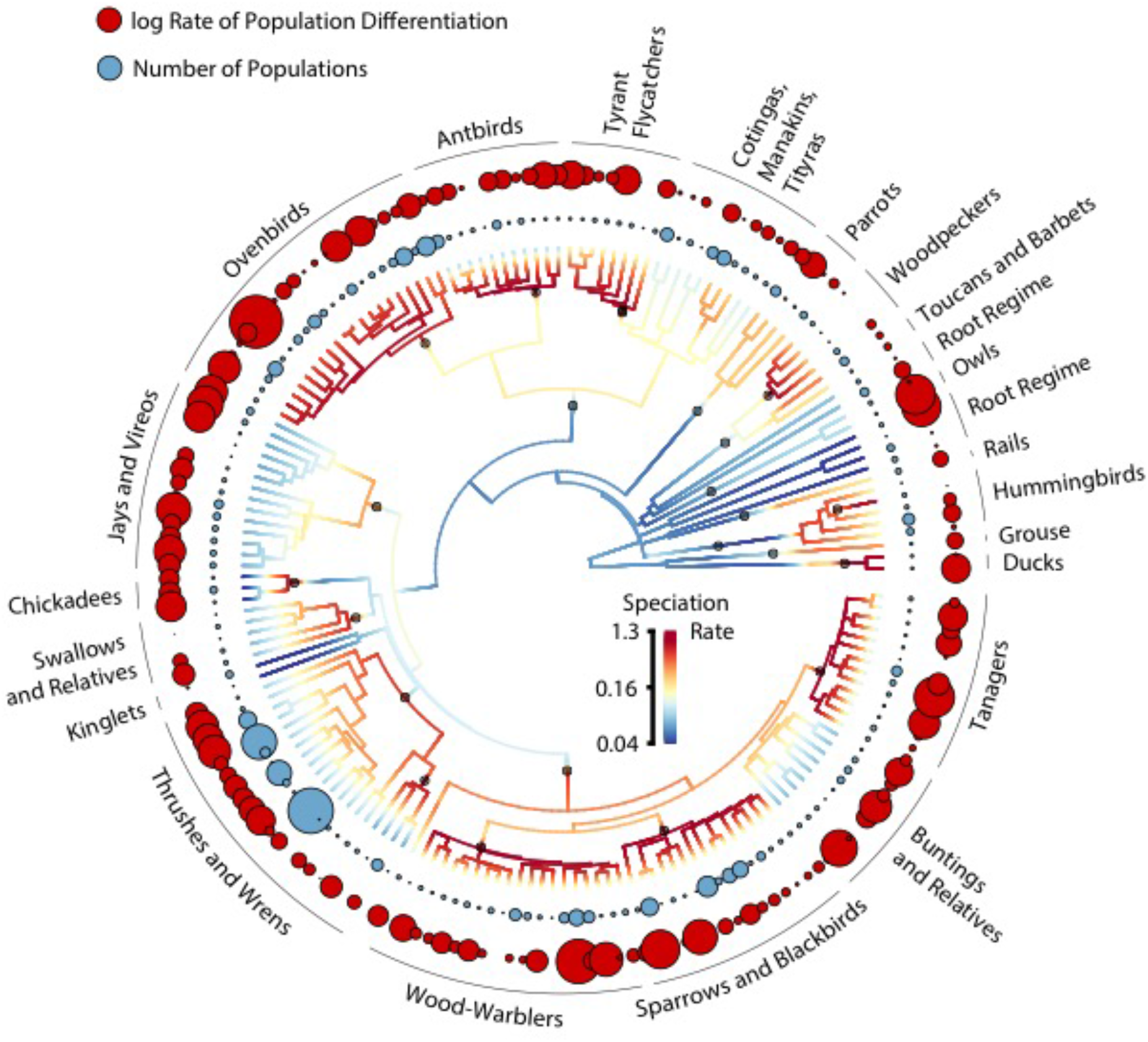
A circular phylogenetic tree of the 176 study species used to estimate rates of population genetic differentiation, colored with a gradient that depicts speciation rates along each branch. The set of 23 macroevolutionary regime shifts with the maximum *a posteriori* probability is plotted on the phylogeny with black circles. The diameter of the blue circles encircling the tree is proportional to the number of populations based on bGMC anlaysis within the adjacent terminal species. The diameter of the red circles at the periphery of the plot is proportional to the log-transformed rate of population differentiation since the crown age of each terminal species.

We estimated macroevolutionary speciation rates along the ancestral lineage leading to each of the 176 species in the population genetic datasets using two methods applied to an existing phylogenetic tree of all bird species (20). First, we computed a simple summary metric of net speciation rate for each tip based on the weighted number of phylogenetic splits between the tip and the root of the phylogeny. This metric was used by Jetz et al. (20) and referred to as the diversification rate (DR) statistic, although it is more tightly related to speciation rate than diversification rate in many cases (21). Speciation rates based on the DR statistic ranged from 0.03 to 3.35 species/My with a median of 0.18 species/My. Second, we used BAMM v.2.5, a Bayesian implementation of a model that jointly estimates the number of distinct evolutionary rate regimes across a phylogenetic tree and the speciation and extinction rates within each of the regimes (22, 23). Speciation rates for tips on the tree were extracted from the marginal distribution of rates for their terminal branches. Based on BAMM analysis, diversification in birds was characterized by 69 statistically distinguishable rate regimes (Fig. S1), 23 of which included the 176 species in our population genetic dataset. The speciation rate across the 176 species varied from 0.04 to 0.72 species/My, with a median of 0.14 species/My (Figs. 2, S2). Importantly, speciation rates inferred using both methods were slower than population differentiation rates. Across all study species, 3.23 times more population differentiation occurred than speciation using the DR statistic, or 4.80 times more using the BAMM speciation rate. These ratios are likely to be conservative because they do not account for population extinction since the crown age of each species. Although most geographic variants are ephemeral and do not persist to become reproductively isolated species, this does not obviate the possibility that variation among lineages in differentiation rate predicts variation in speciation rates.

We tested whether population differentiation rates within species were associated with speciation rates inferred using both BAMM diversification analyses and the number of split summary statistic. We tested for a relationship between population differentiation and speciation rates estimated in BAMM using STRAPP, a trait-dependent diversification test that avoids phylogenetic pseudoreplication while accounting for autocorrelation in evolutionary rates within evolutionary regimes inferred using BAMM (24). We found BAMM speciation rates were positively correlated with population genetic differentiation rates (Spearman’s correlation coefficient [*r*] = 0.250, *P* = 0.021, Fig. 3*A*). We compared population differentiation and speciation rates based on the DR statistic using phylogenetic generalized least squares (PGLS; 25, 26). This test is analogous to that first developed and applied to trait-dependent diversification analyses by Freckleton et al. (27). As with BAMM speciation rate, population differentiation rate predicted speciation rate based on the DR statistic across all study species (PGLS slope = 0.201, *P* < 0.001; Fig. 3*B*). Using simulations, we found the false positive rate, which is problematic in many tests of trait-dependent diversification, was minimal for both the STRAPP (0.022) and PGLS (0.056) analyses relative to a Spearman’s correlation not accounting for covariance (0.568 using BAMM speciation rates, 0.202 using the DR statistic). We found no correlations between the raw number of population clusters and speciation rate (BAMM *r* = 0.104, *P* = 0.313; PGLS slope = -0.077, *P* = 0.111), suggesting that the rate of population genetic differentiation rather than the level of standing differentiation is associated with the rate of speciation. This result provides quantitative evidence supporting the idea that population differentiation within species predicts macroevolutionary dynamics at a large spatial and taxonomic scale.

**Fig. 3.**
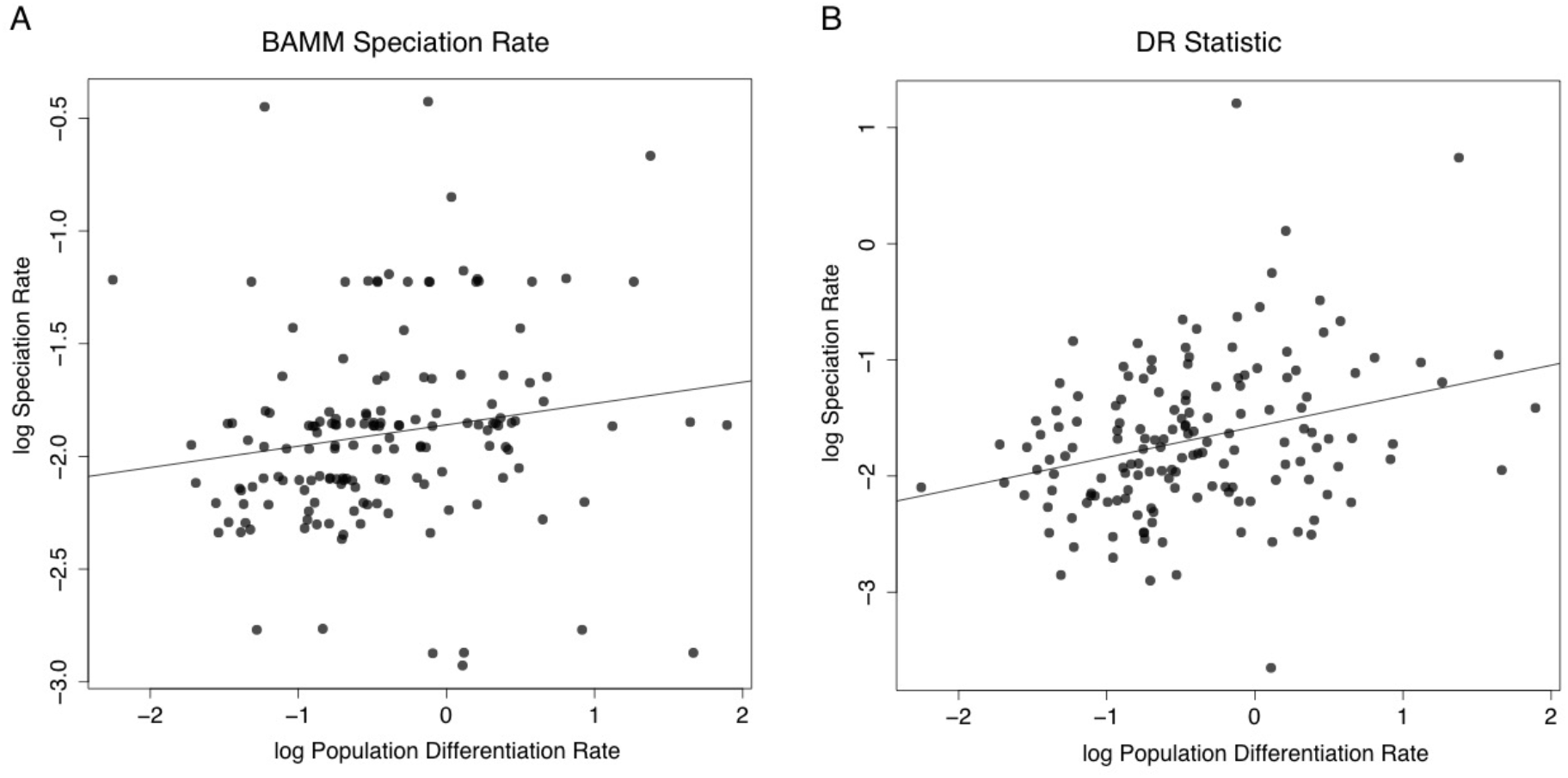
Plots showing the relative population differentiation and speciation rates across all 176 study species. Plots are presented based on speciation rates from (A) BAMM analysis (BAMM correlation coefficient (*r*) = 0.250, *P* = 0.021) and (B) the DR statistic (PGLS slope = 0.201, *P* < 0.001). The trend line for the plot using BAMM speciation rates is based on OLS regression.

A series of tests ensured our results are not dependent on sampling, particular methodological decisions, or statistical artifacts. For brevity, we present results from STRAPP tests of BAMM speciation rates below, but results from PGLS of the DR statistic were similar and are presented in the *SI Materials and Methods*. The positive correlation between the population genetic differentiation rate and the speciation rate was robust to the taxonomy used to circumscribe species for the population-level analysis, with a more finely subdivided taxonomy producing similar results to the primary taxonomy we examined (*r* = 0.210, *P* = 0.014). The correlation was also robust to the use of lower (PP = 0.7; *r =* 0.243, *P =* 0.019) and higher (PP = 0.9; *r =* 0.251, *P =* 0.026) posterior probability thresholds for assigning individuals to population clusters, to whether the population differentiation rate was measured using the stem age rather than crown age of a species (*r* = 0.242, *P* = 0.012), to the random removal of 20% (*r* = 0.238, *P* = 0.027) of samples from the dataset, and to models of population differentiation incorporating moderate (*eps* = 0.45; *r* = 0.199, *P* = 0.043) or high (*eps* = 0.9; *r* = 0.227, *P* = 0.046) extinction rates. Population differentiation rate might be associated with speciation rate if clades with high speciation rates necessarily have shallower crown ages that result in elevated differentiation rates. However, crown age was unrelated to speciation rate (*r* = -0.1134, *P* = 0.2816).

After dividing the species into Tropical (*n* = 101) and Temperate (*n* = 75) groups, we found a strong positive correlation between population differentiation and speciation rates in the tropical species (*r* = 0.409, *P* = 0.005, Fig. 4a), but no correlation in the temperate species (*r* = 0.0358, *P*= 0.7456). The correlation coefficients in tropical and temperate species were more different than expected based on a random permutation test to account for differences in sample sizes between the two groups (*P* = 0.001). Neither population differentiation rate nor speciation rate varied significantly with latitude (Fig. 4A), nor was the average ratio of population differentiation to speciation rate different between temperate and tropical species There was, however, a large disparity across species in the ratio of population differentiation to speciation rates at temperate latitudes, compared to a more peaked distribution in the Tropics (F-test of equal variances F = 2.002, *P* = 0.002; Fig. 4B,C). These results suggest that population differentiation leads to speciation at a relatively predictable rate in the Tropics, but that this rate is less predictable in the Temperate Zone.

**Fig. 4.**
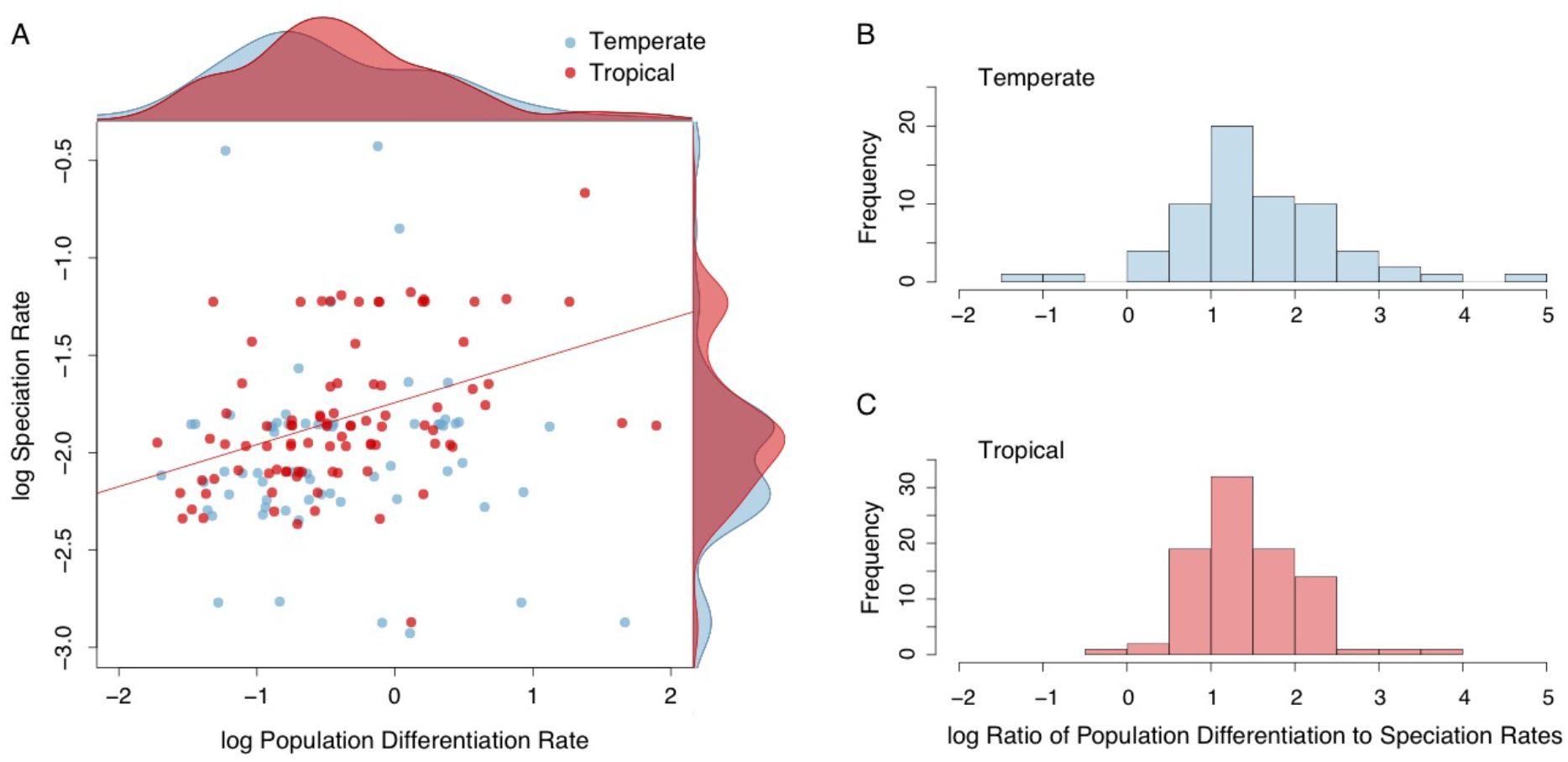
Plots showing differences in relative population differentiation and speciation rates between temperate (*n* = 75) and tropical (*n* = 101) species. (A) Tropical species show a relationship between population differentiation rates and speciation rates (r = 0.409, *P* = 0.005), whereas temperate species do not (*r* = 0.0358, *P* = 0.7456). Kernel density plots showing the relative distributions of rates between tropical and temperate species are plotted opposite the axis of the rate to which they correspond and show that neither differentiation or speciation rates differ noticeably between temperate and tropical species. The ratio of population differentiation rate to speciation rate, however, is more variable in temperate species (B) than tropical species (C). Plots reflect values based on speciation rates estimated using the BAMM speciation rates (see Fig. S3 for plots using the DR statistic).

## Discussion

We found a robust association between population differentiation rate and speciation rate across New World birds, although considerable variance in speciation rate remained unexplained. Given the potential for estimation error in both population differentiation rates and speciation rates, detecting any association was remarkable. This result suggests that the rate of population differentiation within a species can be used, in part, to predict its speciation rate over longer timescales, and vice versa. Statements of causality between rates of population differentiation and speciation would be misleading, however. Population differentiation may be a rate-limiting step in speciation, or both may be related through an unresolved, underlying causal structure involving shared processes that affect rates at both timescales. In either case, our results support an implicit but largely untested assumption of speciation research, that the often-studied processes leading to population differentiation could also be responsible for elevated diversification rates over deep evolutionary time. Our results accord well with prior evidence that taxonomic subspecies richness within species is tied to species richness or speciation rate in their higher taxonomic groups (15, 16) and that some traits predict diversity both within and across species (28). Further support for the link between differentiation and diversification comes from the observation that certain traits thought to lead to population differentiation, such as limited dispersal ability or range fragmentation, predict speciation rate in certain clades (29, 30, 31).

Much of the variance we observed in speciation rates is unexplained by population differentiation. This unexplained variance may be partly due to estimation error in speciation or differentiation rates, but it also leaves room for the possibility that population-level processes in addition to differentiation act as controls on speciation rate. For allopatric speciation to be complete, geographically isolated populations must not only differentiate, but persist until the evolution of reproductive isolation and ecological divergence permit the completion of speciation (6, 13). Variation among species in population persistence, time to reproductive isolation, or time to ecological differentiation therefore may explain some of the variance in speciation rate not attributable to differences in population differentiation rate. However, extinction is notoriously difficult to estimate from molecular phylogenies alone (9, 32), and measuring population persistence for exploration of this potential control using empirical data from extant populations may be equally challenging. Associations between rates of reproductive isolation and speciation rate have been investigated in some groups. The rate of intrinsic postzygotic reproductive isolation does not predict speciation rate across birds (33), but other forms of reproductive isolation are potentially more important in birds (34) and may merit further investigation. Elevated ecological opportunities can be associated with increased speciation rates (35) and there is evidence rates of ecological divergence vary regionally (36), but more data are needed to establish a link between rates of ecological divergence between populations and speciation rate. Regardless, although variation among lineages in rates of population persistence, evolution of reproductive isolation, and ecological divergence may explain some variation in avian speciation rate, they are insufficient to erase the association between population differentiation and speciation observed in our datasets.

Population differentiation predicts speciation rate across all New World birds examined, but the relationship is strongest among tropical species. Tropical species, moreover, show less variability in the rate at which differentiated populations become species compared to temperate species. Because the inverse of the ratio of population differentiation to speciation rates provides an index of population extinction rates, this pattern also indicates that population extinction rates, although not higher in the Temperate Zone, are more variable across temperate species than tropical species. This pattern is consistent with a scenario in which the conversion of population differentiation to new species occurs predictably through time in the Tropics, but is episodic or unpredictable at temperate latitudes. Climatic cycling over the past 420,000 years (37) suggests that major shifts in external environmental conditions may be the dominant driver of speciation rates in those regions, which could dampen the association between population differentiation and speciation at high latitudes. The tighter association between population splitting and speciation rates in the Tropics may be due to the relative environmental stability in that region over recent timescales (38), which might relegate control of speciation rates to the population-level processes occurring constantly within lineages. The latitudinal difference in the correlation between population differentiation and speciation therefore supports hypotheses that invoke greater tropical environmental stability as a cause of the latitudinal diversity gradient (39, 40), and suggests an underlying mechanism in the form of less episodic diversification dynamics resulting from less dramatic climatic shifts.

In conclusion, we suggest that further research on population differentiation is warranted because it captures responses to the same processes that are responsible for organismal diversification generally. We anticipate more and larger comparative, population-level genetic datasets will allow investigation of additional processes responsible for the diversity of organisms worldwide. Furthermore, we expect traits associated with processes that promote population differentiation will prove promising in searches for the attributes of organisms that predispose them to diversify.

## Methods

### Sampling

We examined population genetic data from 176 species from across the New World (*SI Appendix*). We limited our analysis to mainland New World taxa to help control for the area available to each species for accruing allopatric divergence (i.e. differentiation via geographic isolation of populations). Five species whose distributions extend into the Old World were included, but samples from Old World populations were not examined. Species were defined as all non-sympatric monophyletic populations for which we had sampling, regardless of their current treatment by taxonomic authorities. Thus, metrics of population differentiation reflect geographic patterns of diversity among allopatric or parapatric groups, whereas metrics of speciation reflect deeper patterns among potentially sympatric and reproductively isolated groups.

### Controlling for Taxonomic Bias

Despite our attempt at a standardized taxonomy, differences among geographic regions in taxonomic treatment may result in biased results. We alleviated the possibility of taxonomic bias by focusing on rates of differentiation rather than standing levels of differentiation (see below). We expect differentiation rates to be similar in a species regardless of the taxonomic treatment used, because a more inclusive treatment for a given species will generally result in an older species age in addition to more genetic structure. We also investigated the effect of taxonomic treatment on our results by applying a second taxonomy corresponding to the current taxonomy of the American Ornithologist’s Union (AOU) North American (41, 42) and South American (43) checklist committees. In situations where the North and South American committees differed in their treatment, we reverted to the North American committee’s treatment. The AOU taxonomy is more subdivided or “split” (206 species) than the primary taxonomy (176 species), so examination of both provides an index of the impact of the level of taxonomic splitting on results.

### Molecular Data

For 146 species, we relied on previously published population-level mitochondrial datasets of New World birds, restricting our sampling to those datasets containing at least 8 samples (mean = 95) and range-wide sampling. We also gathered data from an additional 30 species, again sampling at least 8 individuals (mean = 111) from across the distribution of each species. We extracted total DNA from tissue samples associated with voucher specimens and used polymerase chain reaction to amplify sequence from the mitochondrial genes NADH dehydrogenase 2 (ND2) or Cytochrome b (cyt *b*) using standard primers. We conducted Sanger sequencing on PCR amplicons. We evaluated how robust our results were to the level of sampling within species by randomly pruning 20% and 40% of the tips of the mitochondrial gene trees estimate from the full dataset, re-estimating the number of bGMYC clusters and rates of population differentiation, and conducting trait-dependent diversification tests.

### Population Divergence Estimation

We estimated mitochondrial gene trees for each species using the Bayesian method implemented in BEAST v.1.7.5 (44). All trees were time-calibrated using an uncorrelated relaxed substitution rate based on published avian mitochondrial rates (*SI Materials and Methods*). We included taxa deemed to be sister to study species based on prior phylogenetic work and we extracted stem and crown age estimates for each species from maximum clade credibility (MCC) trees. We quantified phylogeographic structure using a Bayesian implementation of the General Mixed Yule Coalescent model (bGMYC; *SI Materials and Methods*). We used the MCC tree from BEAST for each bGMYC run. bGMYC provides a posterior probability that two sequences belong to the same genetic species which can be used, along with a probability threshold, to determine the number of clusters present. For the primary analysis we used a posterior probability threshold of 0.8 for clustering, but we also examined higher (0.9) and lower (0.7) thresholds.

To account for the fact that species might differ in the number of bGMYC clusters by virtue of differences in their age, we estimated the rate of bGMYC cluster formation, hereafter the phylogeographic splitting rate. We calculated rates using crown age, the time before present of the first intra-specific divergence event. We calculated rates of bGMYC cluster formation under a pure-birth model using formula (6) from Magallón and Sanderson (45). All rates were calculated using a starting diversity of one despite the use of crown age. Crown age in our study corresponds to the first divergence between mitochondrial haplotypes rather than the first divergence between bGMYC clusters, and thus represents a time point when only one bGMYC cluster was present. We also examined rates of population differentiation using the stem age, although crown age is generally superior to stem age for rate estimation because it is positively correlated with diversity (46), increasing the comparability of rate estimates across species and taxonomic treatments. We did not control for area in our rate estimates, because we expect population differentiation to have equivalent evolutionary importance regardless of the size of the area over which it is distributed.

Because we modeled differentiation at shallow time scales, we might assume that extinction is infrequent and pure-birth (Yule) models provide reasonable estimates of differentiation rate. Jointly estimating speciation and extinction is possible using birth-death models and taking advantage of branch length information in population phylogenies. However, many of our population trees contained so few tips that the likelihood surface for parameter estimation was flat and confidence intervals were very large (T. J. Stadler, pers. comm.). Instead, we examined models that estimated speciation provided different fixed extinction rates. We examined differentiation rates using models with moderate (epsilon = 0.45) and high (eps. = 0.9) constant rates of extinction, in addition to a pure-birth model.

### Speciation Rate Estimation

We used time-calibrated MCC trees from a prior phylogeny of all birds (20) for estimation of speciation rates. Jetz et al. placed species lacking genetic data using taxonomic constraints, but we removed these (leaving 6,670 species) for our analyses to eliminate potential artifacts due to incorrect placement. All study species were represented in the phylogeny by genetic data, and tips in the phylogeny were collapsed in cases in which one of our study species was represented by multiple species in the Jetz et al. taxonomy. We estimated speciation rates on the tree topology based on the Hackett et al. (47) backbone using the model implemented in the program BAMM v.2.5 (22, 23). BAMM was run assuming 67% sampling across the avian tree to account for species without genetic data (48; *SI Materials and Methods*). Speciation rates for a given terminal branch on the tree were extracted from the marginal distribution of rates, which is based on all processes sampled at that branch. We also estimated speciation rates using a simple summary statistic (the “DR statistic”) that reflects the number of splitting events subtending each tip on a phylogenetic tree (20). For this analysis we were unable to analytically account for incomplete sampling, so we used the full avian phylogeny.

### Comparative Analyses

We examined correlations between the population genetic differentiation rate and speciation rates inferred both using the DR statistic and diversification modeling in BAMM. We tested for correlations between log-transformed population differentiation and log-transformed BAMM speciation rates using a semi-parametric trait-dependent diversification test (STRAPP) that detects effects based on replicated associations between trait values and diversification rates estimated using BAMM (24; *SI Materials and Methods*). This test accounts for covariance between species using permutations of trait values amongst species sharing the same evolutionary rate regime. We used PGLS (25, 26) to test for a correlation between log-transformed population differentiation rates and log-transformed DR statistic while accounting for relatedness between species based on phylogenetic distance in the avian tree (20). We tested the rate of false positives (type I error rate) of our STRAPP trait-dependent diversification analysis by simulating trait evolution on our tree under a Brownian motion model (1000 replicates) and conducting the trait-dependent diversification test on the simulated data. We compared the rate of false positives with STRAPP to that based on PGLS using the DR statistic and that based on a simple Spearman’s rank correlation of the population differentiation rates versus BAMM speciation rates.

We conducted comparative analyses on both the full dataset and on datasets containing species from either the Temperate Zone or Tropical Zone. Species were assigned to latitudinal zone based on the latitudinal midpoint of their breeding distribution. We examined partitioning schemes in which south temperate species (*n* = 45) were included either in the tropical partition or the temperate partition. The results were similar in both cases, and we present those with the south temperate species included in the tropical partition, because it resulted in more similar sample sizes between the two partitions (but see *SI Materials and Methods* for results of tests with south temperate species included in the temperate partition).

## Acknowledgments

We thank the field workers and museum staff that obtained and maintain the genetic resources and associated vouchers used in this study, in particular the staff of the Academy of Natural Sciences of Drexel University, American Museum of Natural History, Colección Ornitologica Phelps, Cornell University Museum of Vertebrates, Field Museum of Natural History, Instituto Alexander von Humboldt, Instituto de Ciencias Naturales at the Universidad Nacional de Colombia, Laboratório de Genética e Evolução Molecular de Aves at the Universidade de São Paulo, Louisiana State University Museum of Natural Science, Museo de Historia Natural at the Universidad de los Andes, Museo de Zoología “Alfonso L. Herrera” at the Universidad Nacional Autónoma de Mexico, National Museum of Natural History, University of Kansas Natural History Museum, and University of Washington Burke Museum. We thank A. R. McCune, J. V. Remsen, Jr., S. Singhal, and B. M. Winger for discussion. G. Thomas, A. Crawford, and four anonymous reviewers provided comments that improved the manuscript. Funding was provided by NSF grants DEB-1210556 (to M.G.H. and R.T.B.) and DEB-1146265 (to R.T.B.).

## Footnotes

Author Contributions: M.G.H., G.F.S., B.T.S., and R.T.B. designed the study. M.G.H., G.F.S.,B.T.S., A.M.C., J.T.K., and R.T.B. collected data. M.G.H., G.F.S., and B.T.S. analyzed population-level data. M.G.H. and D.L.R. conducted comparative analyses. All authors contributed to interpretation of the data. M.G.H. wrote the manuscript, and all authors provided comments and revisions.

Genetic data used in this study are available from Genbank (pending acceptance).

Estimates of number of bGMYC clusters, population divergence rates, crown ages, and BAMM speciation rates are avasilable from Dryad (pending acceptance).

Computer scripts used to process data are available from Github (pending acceptance).

## Supporting Information

### SI Materials and Methods

#### Sampling and Taxonomy

Our species sampling was determined by the species for which datasets were currently available or under construction in our labs. We restricted sampling to the mainland New World because New World birds are much better represented in existing genetic resources collections and genetic datasets than Old World species, which are particularly poorly sampled in tropical areas (1). In only a few cases (*Anas strepera, Calidris ptilocnemis, Corvus corax, Hirundo rustica,* and *Pinicola enucleator*) do study species contain Old World populations, which were not sampled for our analyses. Several of these species (the *Anas, Calidris*, and *Pinicola*) are important as the only or one of few representatives of their clade with phylogeographic data available. The phylogenetic dataset used to estimate speciation rates was worldwide in coverage. This strategy allowed us to maintain consistency between datasets – both the population genetic datasets (with the five exceptions above) and the phylogenetic dataset included all subtending lineages rather than a potentially biased subsample. Including all species in the phylogeny, combined with an analytical correction for taxa lacking genetic data, allowed us to estimate absolute speciation rates from each lineage for direct comparison with population differentiation rates.

#### Population Differentiation Rate Estimation

Mitochondrial gene trees were time-calibrated using an uncorrelated relaxed substitution rate based on published avian mitochondrial rates of 0.0125 substitutions/site/My for ND2 and ATPASE6, 7, & 8 (2) and 0.0105s s/s/My for cyt *b* (3). For the gene COI we used the same rate as cyt *b* because the loci mutate at similar rates (2). For the clock rate parameter we specified a lognormal distribution on the prior with the mean set to the above-mentioned mutation rates and a standard deviation of 0.1. We note that these substitution rates may differ from the rates represented in the phylogenetic data. Because substitution rates differ between short and long timescales (4), however, using rates that have been widely applied and tested on intraspecific timescales is the best option in this case. We expect any bias in crown and stem ages from the population-level data relative to branching times from the phylogenetic data to be minimal and have minimal impact on population differentiation rate estimates. Moreover, because we apply the same calibrations for all study species, this bias would be equivalent across species and therefore would not impact correlations between population differentiation and speciation rates.

We estimated time-calibrated gene trees in BEAST using a coalescent-constant-size tree prior and the best-fit nucleotide substitution model as determined in MEGA6 (5). We ran each analysis for 50 million generations sampling every 2,500 generations, performed multiple independent runs for validation, and assessed Markov chain Monte Carlo (MCMC) convergence and determined burn-in by examining ESS values and likelihood plots in Tracer v.1.5 (6). For some datasets that did not achieve high ESS values after 50 million generations, we included additional generations until the results were stable. Maximum clade credibility (MCC) trees were estimated from the posterior distribution of trees for each species using TreeAnnotator (7).

Units delimited using the GMYC model are typically regarded as species. In birds, however, more stringent criteria, involving metrics of reproductive isolation or phenotypic divergence, are typically applied for species delimitation (8, 9). Non-interbreeding populations failing to meet this criteria, such as many of the units delimited in this study, are often assigned subspecies status. Many of the units in this study may even fail to meet some researchers’ criteria for subspecies status, indeed many have not been elevated to subspecies despite prior publications based on the same genetic data examined here. As a result, GMYC units in birds are perhaps better treated as genetically differentiated populations, and we follow this philosophy here. Regardless of their taxonomic status, however, GMYC clusters represent more finely resolved and more recently diverged groups relative to the terminal taxa in the avian phylogeny we examined, and thus are appropriate for comparing divergence rates between recent and deep timescales, the fundamental goal of this study. An alternate method for identifying geographic variants would have been to use named subspecies, but naming practices are far from standardized and many cryptic taxa in poorly studied groups would be missed (10).

bGMYC determines the number of genetic species by estimating the number of clusters within which splits in the gene tree fit a coalescent model rather than a model of interspecific diversification (Yule model). We ran the program for 250,000 generations using the single.phy function and discarded the first 15,000 generations as burn-in. We ran each analysis multiple times for validation, and assessed MCMC diagnostics by examining likelihood plots in Tracer.

We corrected for differences in the age of species by calculating the rate at which bGMYC clusters formed since the species’ crown age. The area across which a species is distributed might also predict its level of differentiation, but we found area was not strongly correlated with the number of differentiated populations (R^2^ = 0.014, *P* = 0.065), certainly much less so than age(R^2^ = 0.298, *P* ≪ 0.001). This suggests that population differentiation has similar evolutionary potential regardless of the size of the area across which it occurs, and we therefore do not control for area in any subsequent analyses.

#### Speciation Rate Estimation

The software program BAMM uses reversible-jump MCMC to examine models differing in the number of time-varying diversification processes present across the phylogeny. Each process includes a time-varying speciation term and a time-invariant extinction rate. In the BAMM run using the primary taxonomy, multiple tips from within the same species were collapsed so as to avoid overlap in the data used for estimation of speciation and population differentiation rates. We ran BAMM using a model allowing for variable rates for at least 350 million generations in both the split and primary analyses, completing multiple runs with the same settings for validation. We sampled every 200,000 generations and discarded 10% of the sample as burn-in. Marginal distributions of speciation rates at the tips of the tree represent estimates of present-day speciation rates for those taxa.

#### Comparative Analyses

We first used STRAPP, which computes the correlation between character states at the tips of the tree and their corresponding diversification rates, and assesses significance by permuting speciation rates among regimes estimated in BAMM. Parametric uncertainty in diversification rates is accommodated by conducting tests across the posterior distribution of rates inferred using BAMM. The permutation test is used to control for the covariance among species from the same macroevolutionary rate regime, thereby explicitly incorporating covariance among replicates with shared history and macroevolutionary dynamics. All tests presented are two-tailed tests, examining the alternative hypothesis that there is a correlation between population differentiation and speciation rates. One-tailed tests, in which the alternative hypothesis is the presence of a positive correlation between population differentiation and speciation rates, resulted in greater significance values than the two-tailed tests presented.

The primary two-tailed test presented in the main text, for example, resulted in a significance level of *P* = 0.021, whereas the equivalent one-tailed test resulted in a level of *P* = 0.007. We used the two-tailed test as a more conservative index of trait-dependent diversification.

We also conducted comparative analyses of the number of splits summary statistic of speciation rate using phylogenetic generalized least squares (PGLS) in the caper package (11) in R. PGLS analysis using the number of splits summary statistic produced similar results to STRAPP overall. In addition to the results presented in the main text, PGLS analysis using the split taxonomy resulted in a correlation similar to that using the primary taxonomy (PGLS slope = 0.182, *P* = 0.003), as did the use of lower (0.7;; PGLS slope = 0.215, *P* < 0.001) and higher (0.9; PGLS slope = 0.236, *P* < 0.001) posterior probability thresholds for assigning individuals to population clusters, to whether the population differentiation rate was measured using the stem age rather than crown age of a species (PGLS slope = 0.342, *P* < 0.001), to the random removal of 20% (PGLS slope = 0.202, *P* < 0.001) and 40% (PGLS slope = 0.189, *P* < 0.001) of samples from the dataset, and to models of population differentiation incorporating moderate (*eps* = 0.45; PGLS slope = 0.183, *P* < 0.001) or high (*eps* = 0.9; PGLS slope = 0.142, *P* = 0.002) extinction rates. PGLS revealed a correlation between differentiation and speciation rates in the partition containing only tropical species (PGLS slope = 0.215, *P* = 0.002), but no correlation in temperate species (PGLS slope = 0.150, *P* = 0.087). The correlation coefficients in tropical and temperate species were more different than expected based on a random permutation test to account for differences in sample sizes between the two groups (*P =* 0.012). The disparity in the species-specific ratios of population differentiation to speciation rates at temperate latitudes was greater than in the Tropics (F-test of equal variances F = 2.094, P = 0.001).

When species were partitioned into temperate and tropical for the main analysis, south temperate species were included in the tropical partition because most are members of largely tropical groups and we therefore expect them to exhibit long-term evolutionary dynamics similar to the tropical species, and because it resulted in more equivalent sample sizes between partitions.Including these species in the temperate partition, however, produced similar results, with no or a weak correlation observed between population differentiation in the temperate zone (BAMM *r* = 0.149, *P* = 0.179; PGLS slope = 0.214, *P* = 0.001) and a strong correlation in the Tropics (BAMM *r* = 0.458, *P* = 0.007; PGLS slope = 0.332, *P* = 0.002).

## SI Figures

**Fig. S1.**
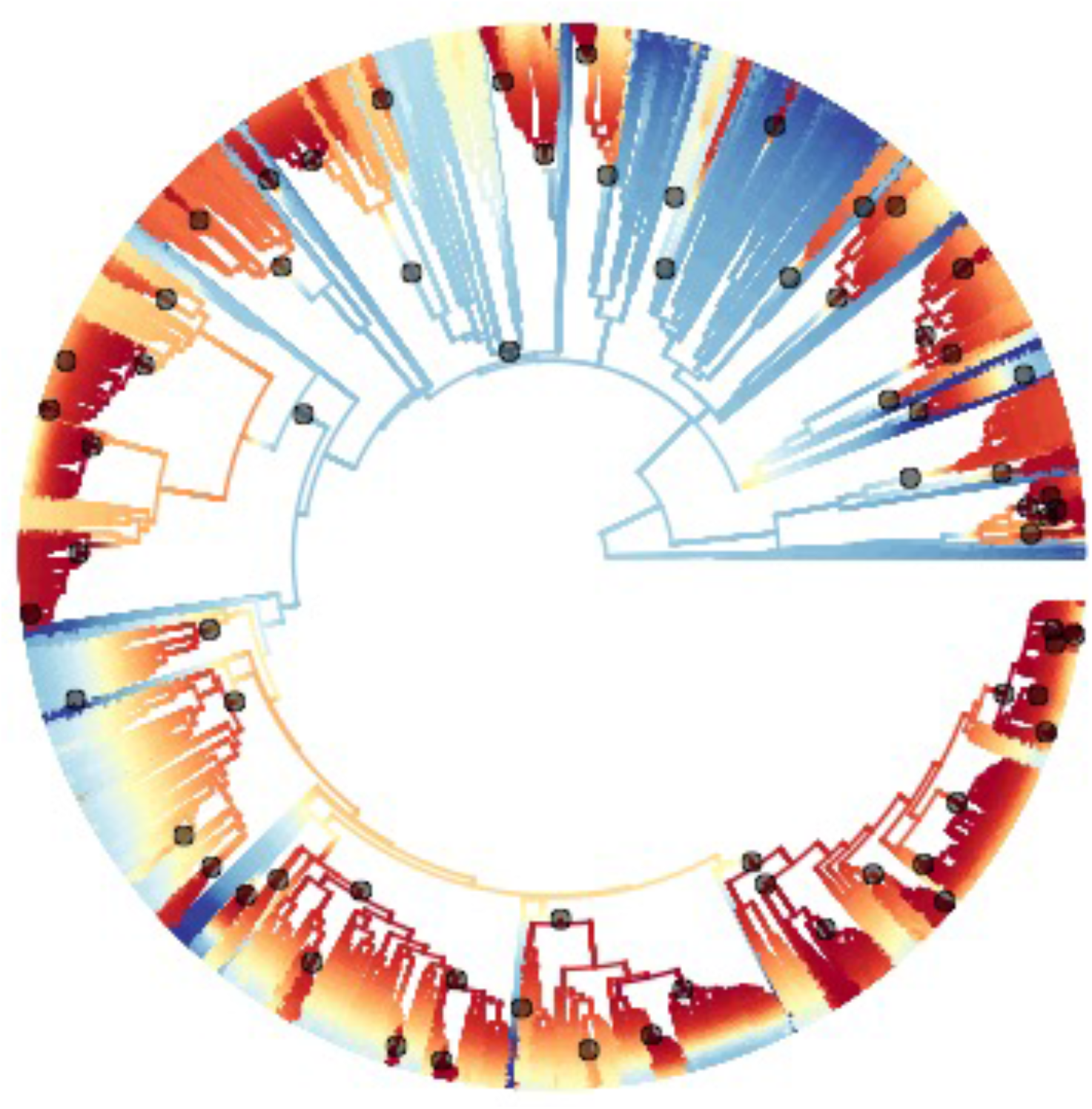
A phylorate plot showing speciation rates and all 69 macroevolutionary regime shifts across the tree of all 6,670 birds with genetic data based on the Jetz *et al.* phylogeny.

**Fig. S2.**
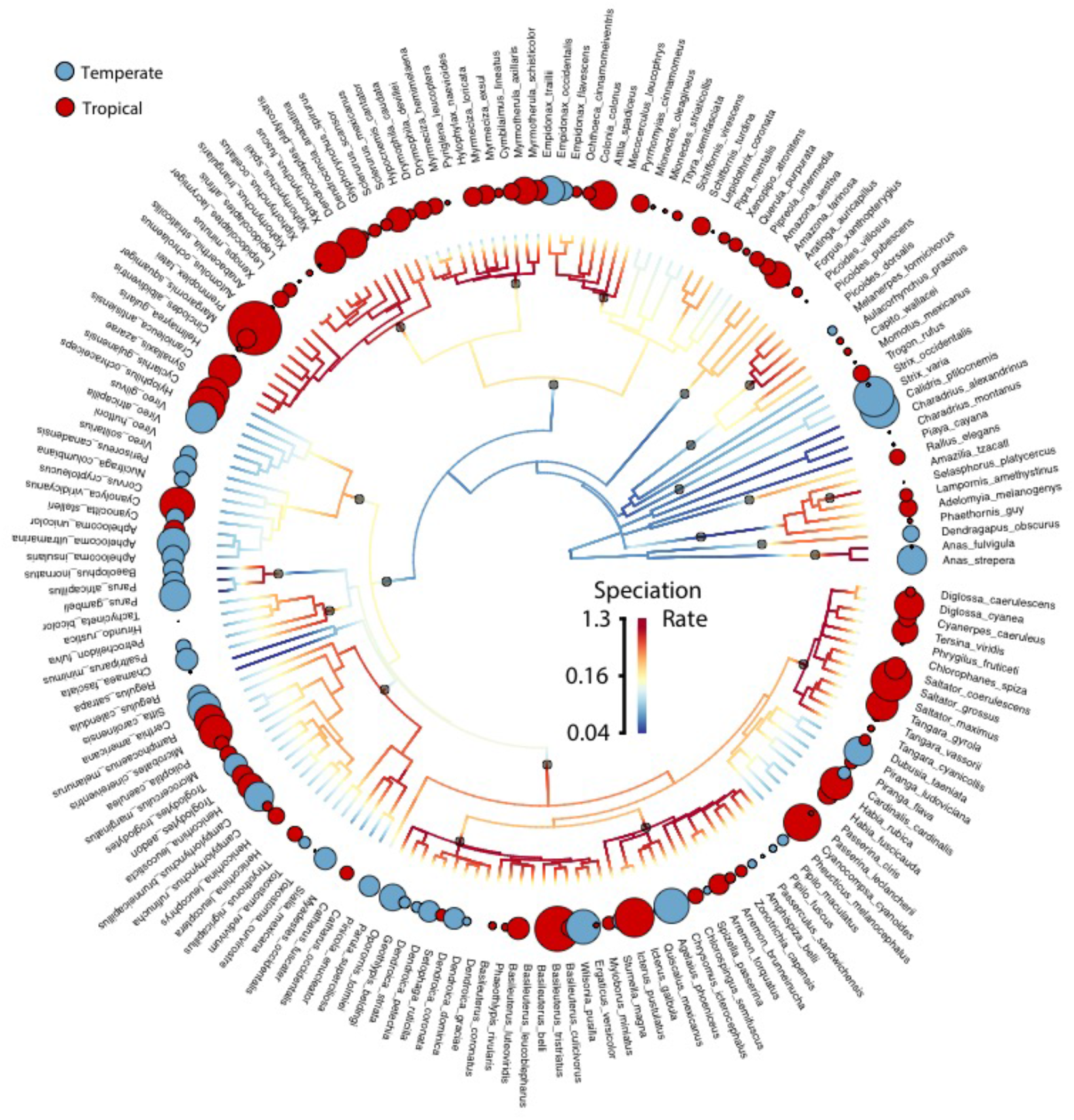
A phylorate plot of the study species similar to that from Fig. 2 in the main text. Colors indicate mean posterior speciation rates along branches. Circles representing log-transformed population differentiation rates of the adjacent tips and are color-coded according to whether each species is tropical or temperate. Species names are listed adjacent to the circle representing their differentiation rate. For species containing multiple AOU species names, the name shown was selected randomly from the name set.

**Fig. S3.**
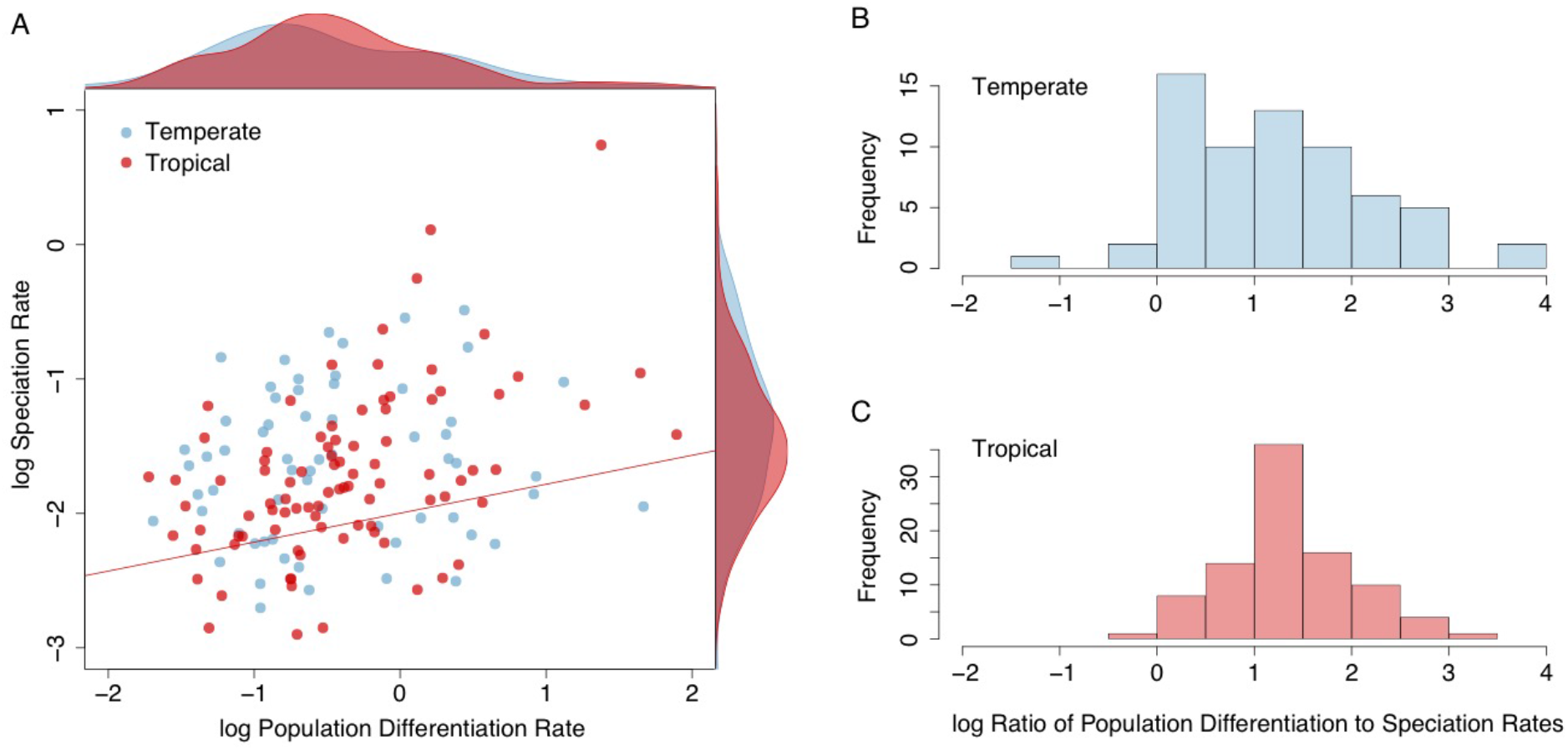
Plots showing differences in relative population differentiation and speciation rates between temperate and tropical species using the DR statistic. (A) Tropical species show a relationship between population differentiation rates and speciation rates (0.215, p = 0.002), whereas tropical species do not. Kernel density plots showing the relative distributions of rates between tropical and temperate species are plotted opposite the axis of the rate to which they correspond and show that neither differentiation or speciation rates differ noticeably between temperate and tropical species. The ratio of population differentiation rate to speciation rate, however, is more variable in temperate species (B) than tropical species (C).

## SI Appendix

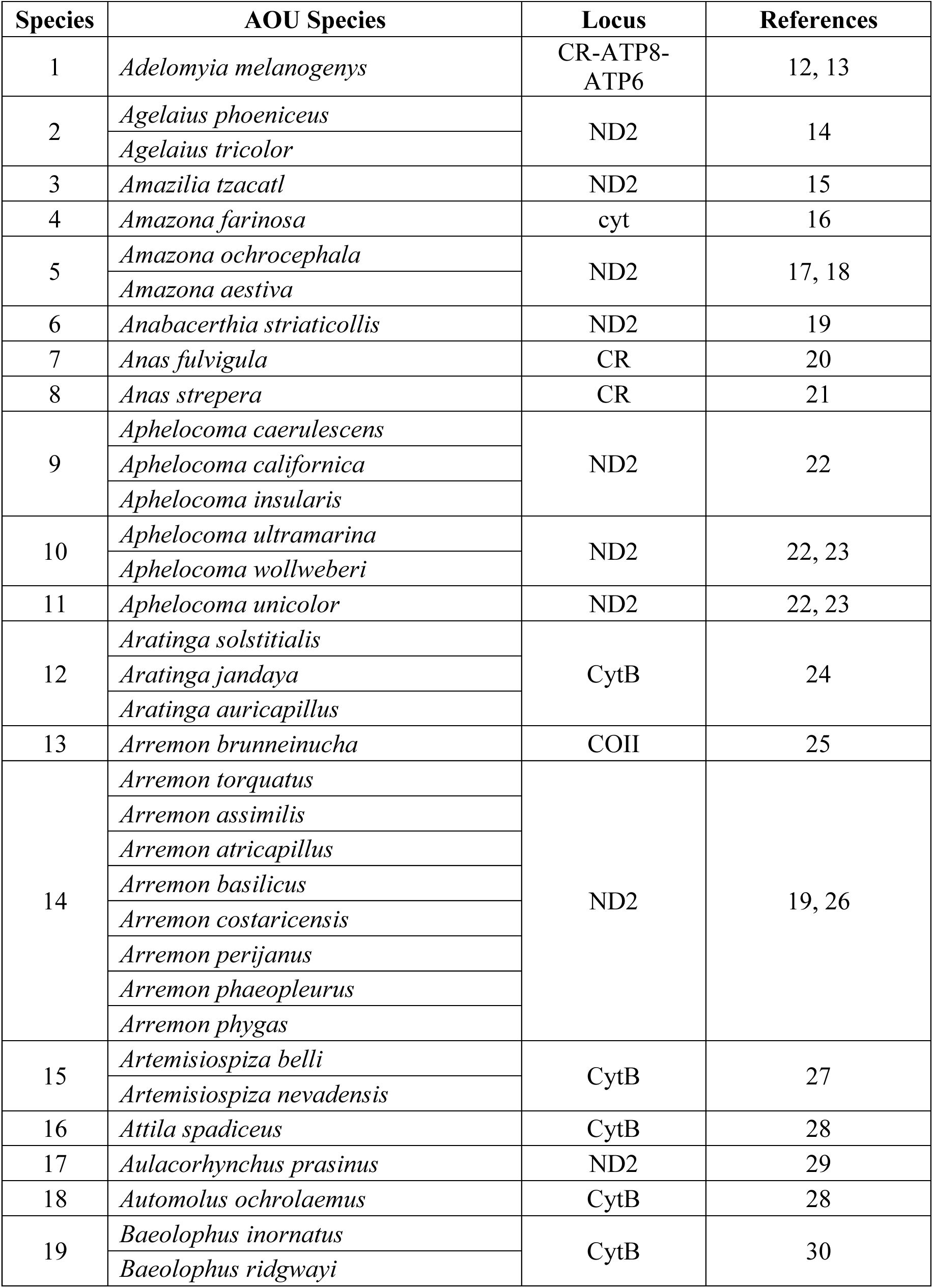
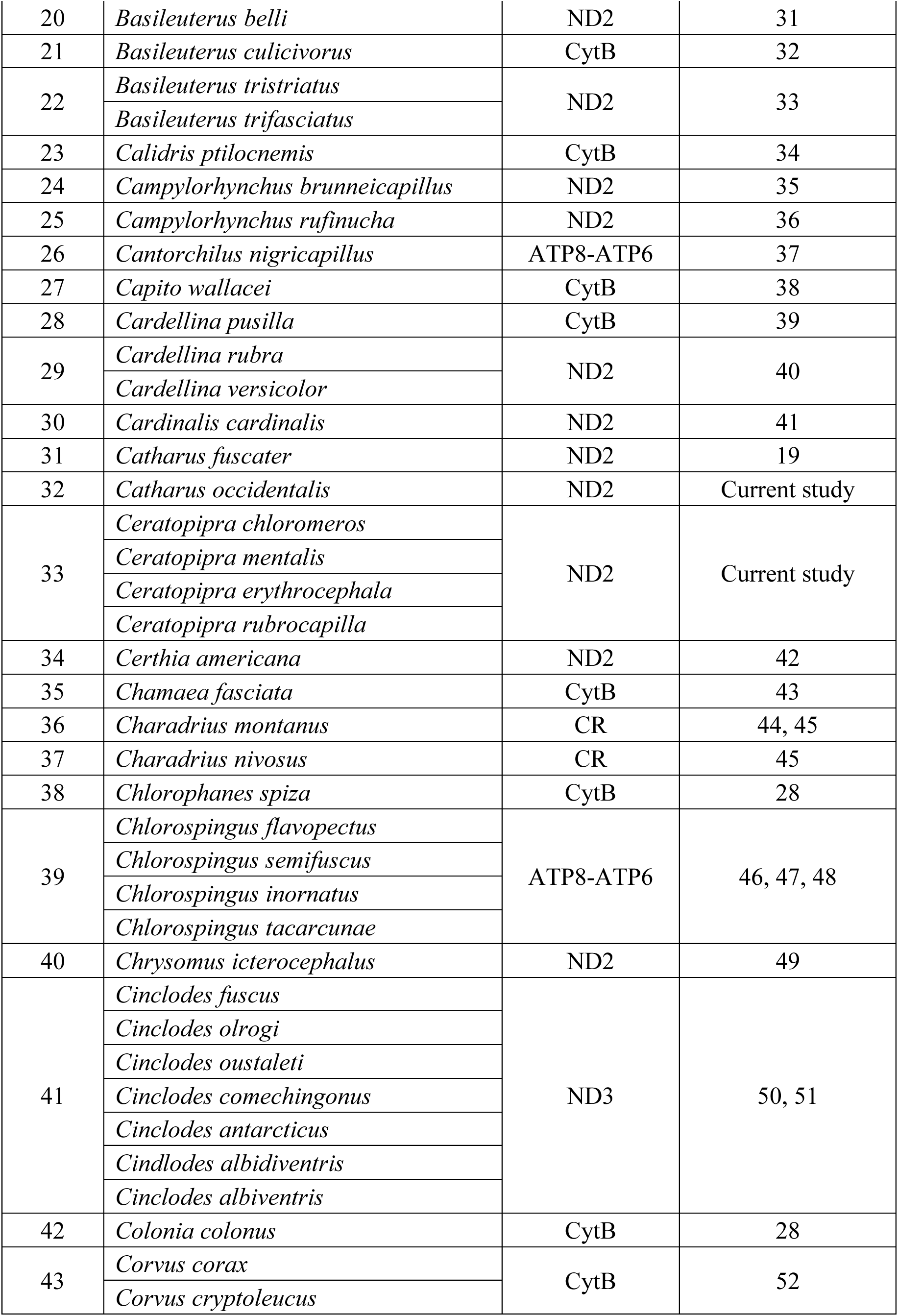
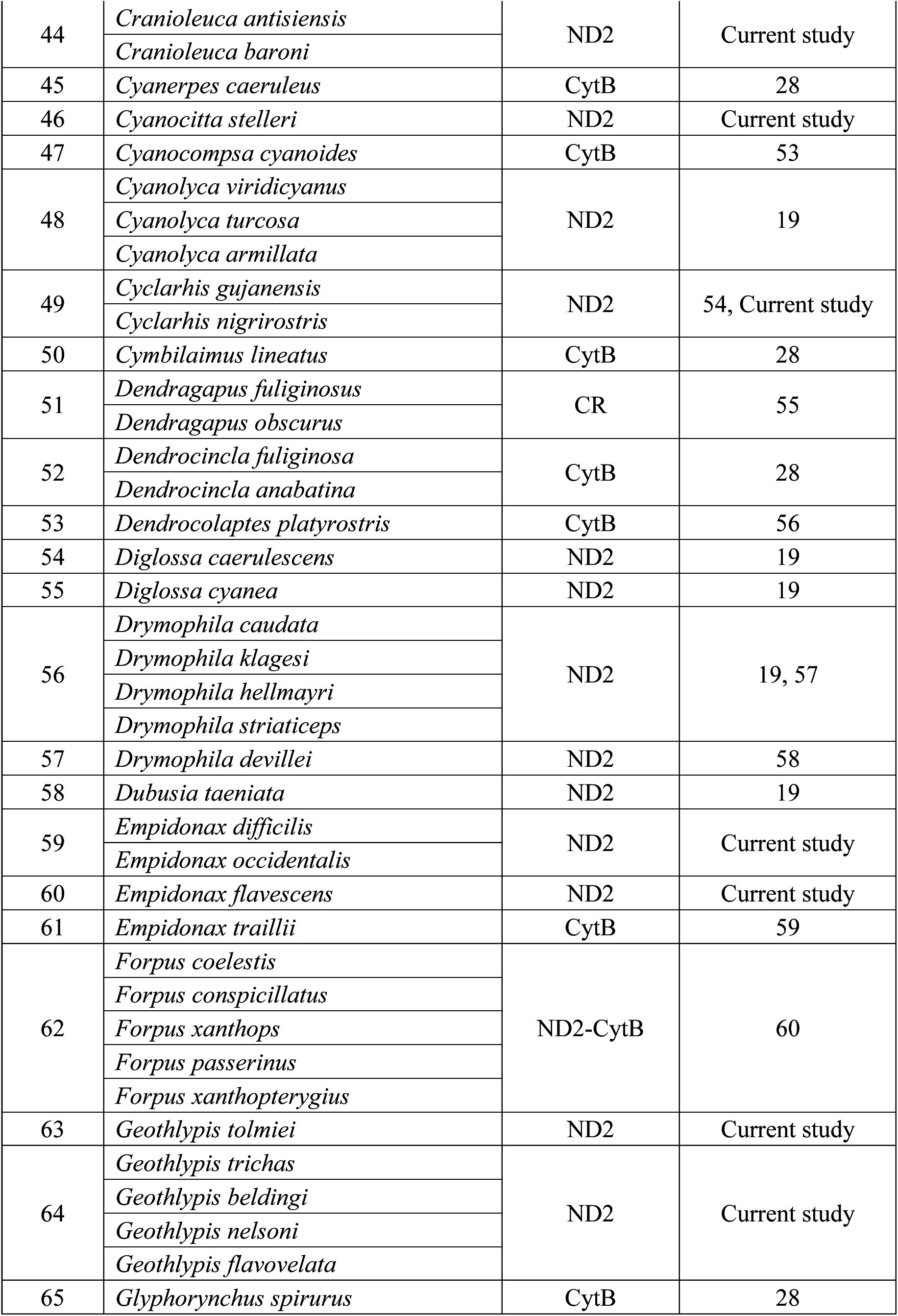
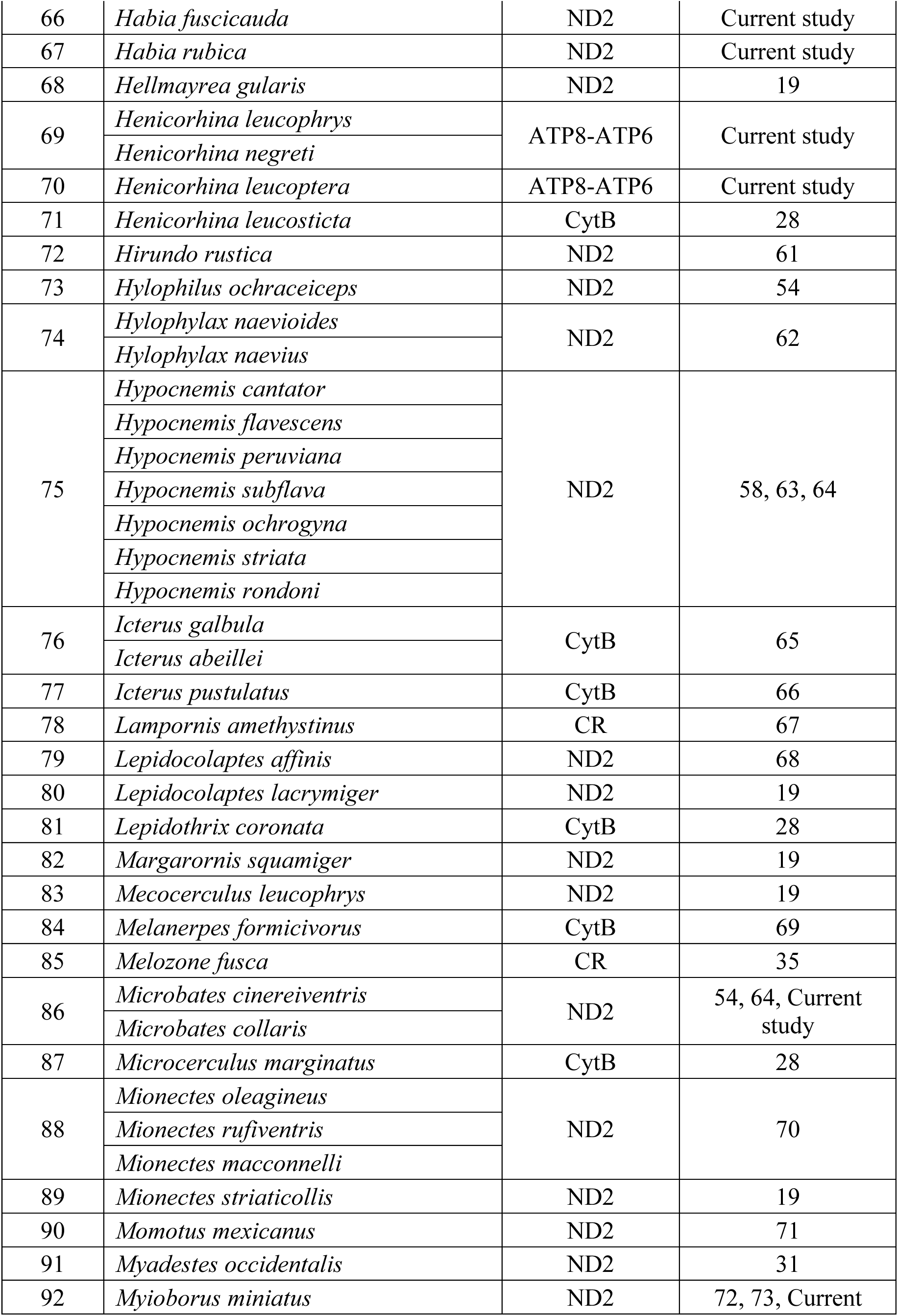
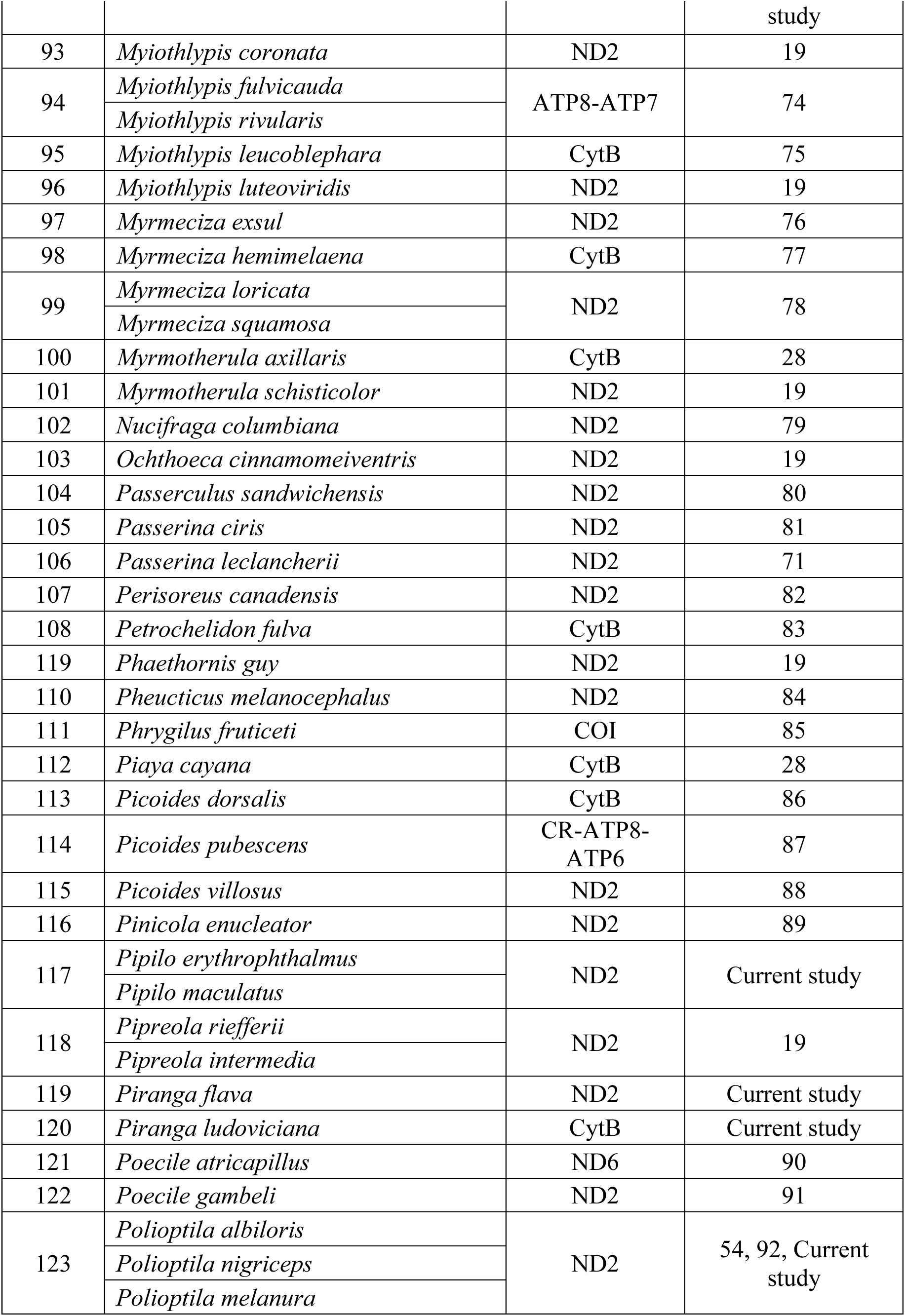
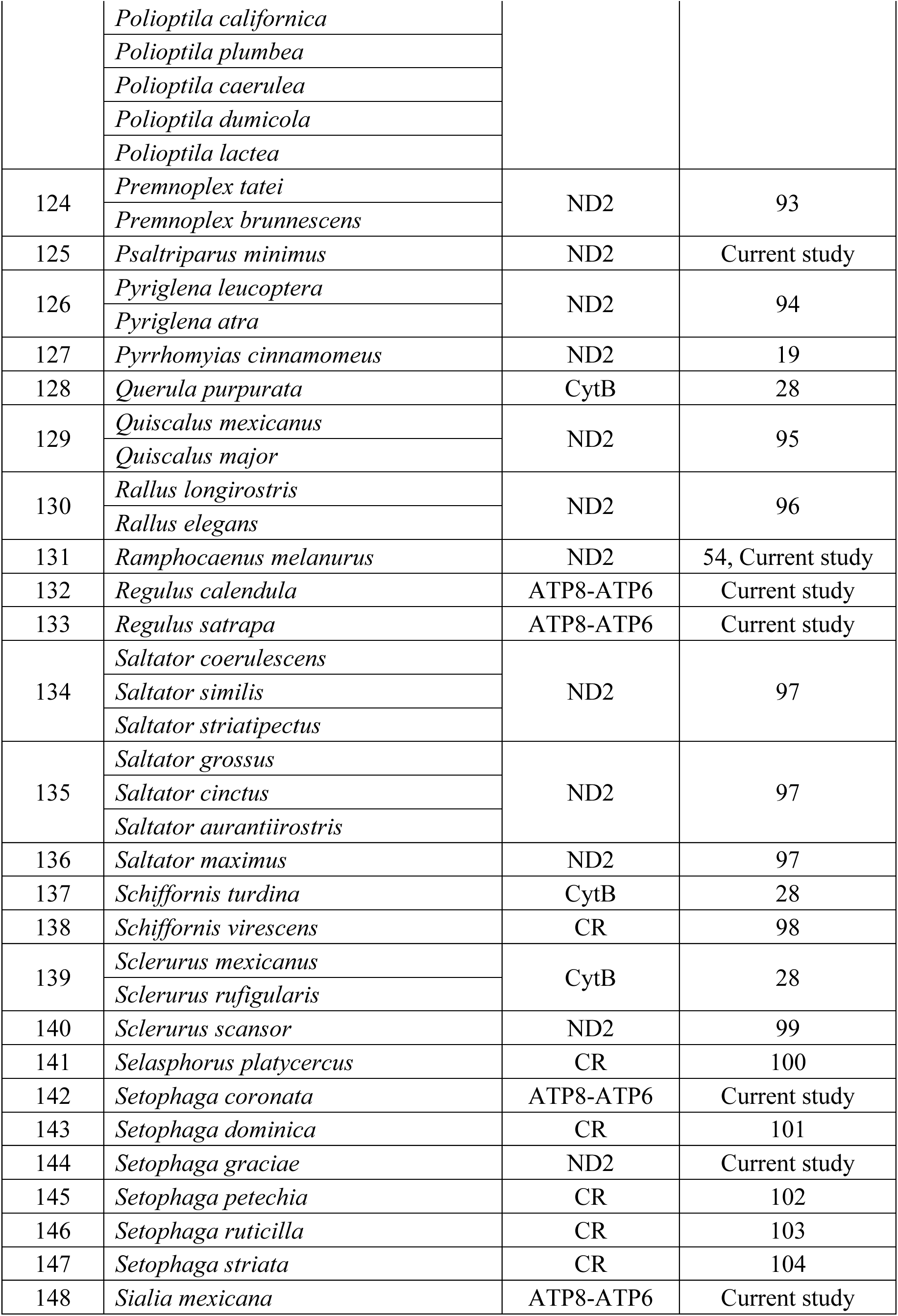
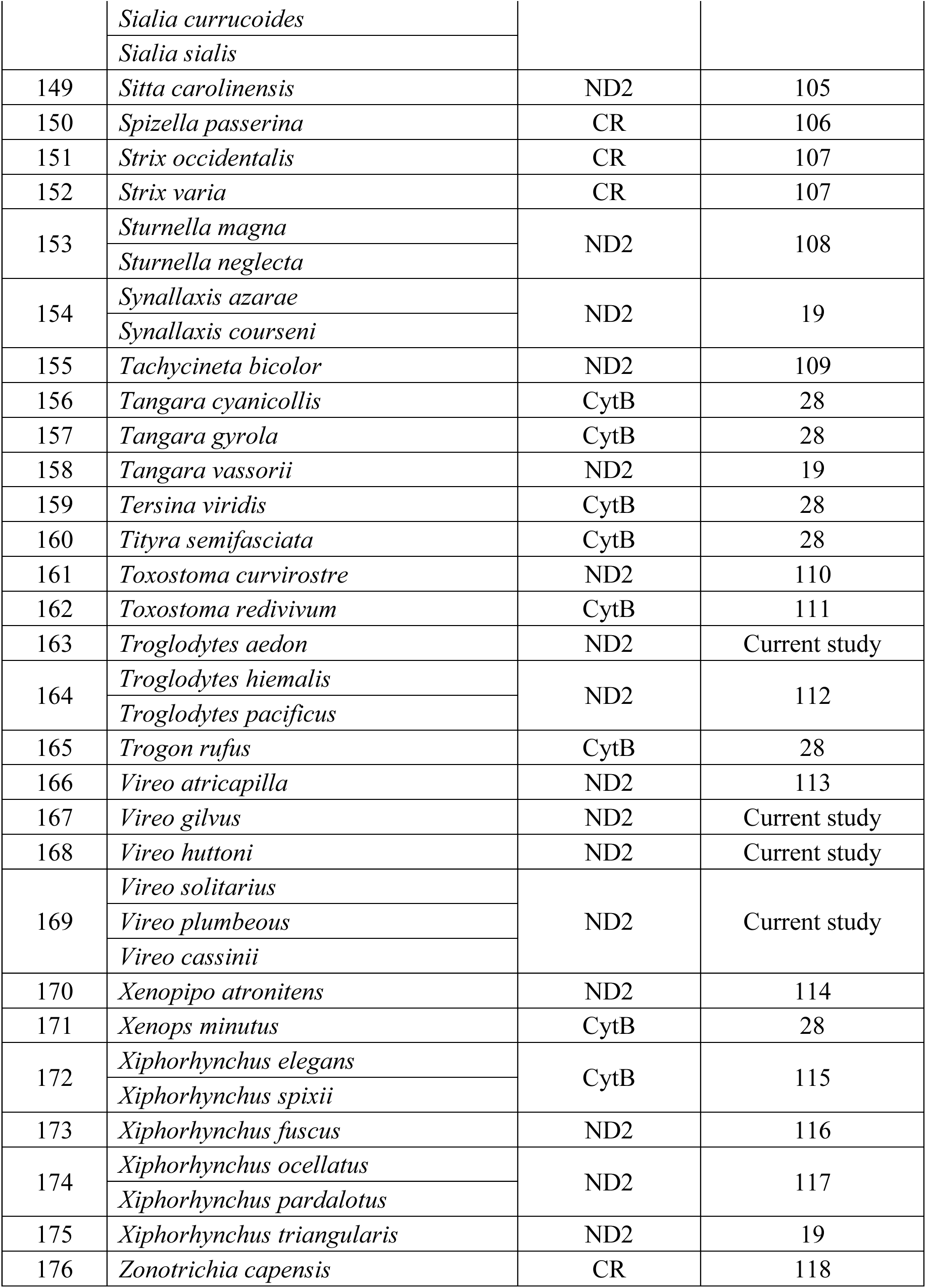

## References

1. Hoorn C, et al. (2010) Amazonia through time: Andean uplift, climate change, landscape evolution, and biodiversity. Science 330:927–931.

2. Wade L (2010) Cradle of life. Science 350:496–501.

3. Smith BT, et al. (2014) The drivers of tropical speciation. Nature 515:406–409.

4. Barreiro LB, Laval G, Quach H, Patin E, Quintana-Murci L (2008) Natural selection has driven population differentiation in moden humans. Nature Genetics 40:340–345.

5. Price TD, et al. (2014) Niche filling slows the diversification of Himalayan songbirds. Nature 509:222–225.

6. Mayr E (1963) Animal Species and Evolution (Belknap Press, Cambridge, MA).

7. Stanley SM (1979) Macroevolution: Pattern and Process (W. H. Freeman and Co., San Francisco).

8. Rosenblum EB, et al. (2012) Goldilocks meets Santa Rosalia: An ephemeral speciation model explains patterns of diversification across time scales. Evol. Biol. 39:255–261.

9. Dynesius M, Jansson R (2014) Persistence of within-species lineages: a neglected control of speciation rates. Evolution 68:923–934.

10. Mayr E (1947) Ecological factors in speciation. Evolution 1:263–288.

11. Dobzhansky T (1937) Genetics and the Origin of Species (Columbia University Press, New York).

12. Mayr E (1942) Systematics and the Origin of Species from the Viewpoint of a Zoologist (Columbia University Press, New York).

13. Allmon WD (1992) A causal analysis of stages in allopatric speciation. Oxford Surv. Evol. Biol. 8:219–257.

14. Barraclough TG, Nee S (2001) Phylogenetics and speciation. Trends Ecol. Evol. 16:391–399.

15. Haskell DG, Adhikari A (2009) Darwin’s manufactory hypothesis is confirmed and predicts extinction risk of extant birds. PLoS One 4:e5460.

16. Phillimore AB (2010) Subspecies origination and extinction in birds. Ornith. Monogr. 67:42–53.

17. Kisel Y, et al. (2012) Testing the link between population genetic differentiation and clade diversification in Costa Rican orchids. Evolution 66:3035–3052.

18. Pons J, et al. (2006) Sequence-based species delimitation for the DNA taxonomy of undescribed insects. Syst. Biol. 55:595–609.

19. Reid NM, Carstens BC (2012) Phylogenetic estimation error can decrease the accuracy of species delimitation: a Bayesian implementation of the general mixed Yule-coalescent model. BMC Evol. Biol. 12:196.

20. Jetz W, Thomas GH, Joy JB, Hartmann K, Mooers, AO (2012) The global diversity of birds in space and time. Nature 491:444–448.

21. Belmaker J, Jetz W (2015) Relative roles of ecological and energetic constraints, diversification rates, and region history on global species richness gradients. Ecology Letters 18:563–571.

22. Rabosky DL (2014) Automatic detection of key innovations, rate shifts, and diversity-dependence on phylogenetic trees. PLoS One 9:e89543.

23. Mitchell JS, Rabosky DL (2016) Bayesian model selection with BAMM: Effects of the model prior on the inferred number of diversification shifts. Methods Ecol. Evol. doi: doi:10.1111/2041-210X.12626.

24. Rabosky DL, Huang H (2015) A robust semi-parametric test for detecting trait-dependent diversification. Syst. Biol. 65:181–193.

25. Grafen A (1989) The phylogenetic regression. Philos. Trans. R. Soc. London B 326:119–157.

26. Martins EP, Hansen TF (1997) Phylogenies and the comparative method: a general approach to incorporating phylogenetic information into the analysis of interspecific data. Am. Nat. 149:646–667.

27. Freckleton RP, Phillimore AB, Pagel M (2008) Relating traits to diversification: A simple test. Am. Nat. 172:102–115.

28. Riginos C, et al. (2014) Dispersal capacity predicts both population genetic structure and species richness in reef fishes. Am. Nat. 184:52–64.

29. Jablonski D (1986) Larval ecology and macroevolution in marine invertebrates. Bull. Mar. Sci. 39:565–587.

30. Owens IPF, Bennett PM, Harvey PH (1999) Species richness among birds: body size, life history, sexual selection, or ecology? Proc. R. Soc. B 266:933–939.

31. Claramunt S, Derryberry EP, Remsen JV, Brumfield RT (2012) High dispersal ability inhibits speciation in a continental radiation of passerine birds. Proc. R. Soc. B 279:1567–1574.

32. Rabosky DL (2010) Extinction rates should not be estimated from molecular phylogenies. Evolution 64:1816–1824.

33. Rabosky DL, Matute DR (2013) Macroevolutionary speciation rates are decoupled from the evolution of intrinsic reproductive isolation in Drosophila and birds. Proc. Nat. Acad. USA 110:15354–15359.

34. Price TD (2008) Speciation in Birds (Roberts and Co., Greenwood Village, CO).

35. Schluter D (2000) The Ecology of Adaptive Radiation (Oxford University Press, Oxford).

36. Lawson AM, Weir JT (2014) Latitudinal gradients in climatic-niche evolution accelerate trait evolution at high latitudes. Ecol. Lett. 17:1427–1436.

37. Petit JR, et al. (1999) Climate and atmospheric history of the past 420,000 years from the Vostok ice core, Antarctica. Nature 399:429–436.

38. Thomson LG, Mosley-Thomson E, Henderson KA (2000) Ice-core palaeoclimate records in tropical South America since the Last Glacial Maximum. J. Quat. Sci. 15:377–394.

39. Hewitt G (2000) The genetic legacy of the Quaternary ice ages. Nature 405:907–913.

40. Weir JT, Schluter D (2007) The latitudinal gradient in recent speciation and extinction rates of birds and mammals. Science 315:1574–1576.

41. American Ornithologists’ Union or AOU (1998) Check-list of North American Birds, 7^th^ Edition (American Ornithologists’ Union, Washington, D. C.).

42. Chesser, RT et al. (2013) Fifth-fourth supplement to the American Ornithologists’ Union Check-List of North American Birds. Auk 130:558–571.

43. Remsen JV, et al. (2014) A classification of the bird species of South America. AOU South American Classification Committee. http://www.museum.lsu.edu/˜Remsen/SACCBaseline.htm.

44. Drummond AJ, Suchard MA, Xie D, Rambaut A (2012) Bayesian phylogenetics with BEAUti and the BEAST 1.7. Mol. Bio. Evol. 29:1969–1973.

45. Magallón S, Sanderson MJ (2001) Absolute diversification rates in angiosperm clades. Evolution 55:1762–1780.

46. Stadler T, Rabosky DL, Ricklefs RE, Bokma F (2014) On age and species richness in higher taxa. Am. Nat. 184:447–455.

47. Hackett SJ, et al. (2008) A phylogenomic study of birds reveals their evolutionary history. Science 320:1763–1768.

48. Rabosky DL, Title PO, Huang H (2015) Minimal effects of latitude on present-day speciation rates in New World birds. Proc. R. Soc. B 282:20142889.

## References

1. Reddy S (2014) What’s missing from avian global diversification analyses? Mol. Phylo. Evol. 77:159–165.

2. Smith BT, Klicka J (2010) The profound influence of the Late Pliocene Panamanian uplift on the exchange, diversification, and distribution of New World birds. Ecography 33:333–342.

3. Weir JT, Schluter D (2008) Calibrating the avian molecular clock. Molecular Ecology 17:2321–2328.

4. Ho SYW, et al. (2011) Time-dependent rates of molecular evolution. Mol. Ecol. 20:3087–3101.

5. Tamura K, Stecher G, Peterson D, Filipski A, Kumar S (2013) MEGA6: Molecular Evolutionary Genetics Analysis Version 6.0. Mol. Biol. Evol. 30:2725–2729.

6. Rambaut A, Drummond AJ (2007) Tracer v.1.5. http://tree.bio.ed.ac.uk/software/tracer.

7. Rambaut A, Drummond A (2008) TreeAnnotator v1.4.8. http://beast.bio.ed.ac.uk/TreeAnnotator.

8. McKitrick MC, Zink RM (1988) Species concepts in ornithology. Condor 90:1–14.

9. Gill FB (2014) Species taxonomy of birds: Which null hypothesis? Auk 131:150–161.

10. Zink RM (2004) The role of subspecies in obscuring avian biological diversity and misleading conservation policy. Proc. R. Soc. B 271:561–564.

11. Orme D, et al. (2013) caper: Comparative analyses of phylogenetics and evolution in R. R package version 0.5.2. http://CRAN.R-project.org/package=caper.

12. Chavez JA, Pollinger JP, Smith TB, LeBuhn G (2007) The role of geography and ecology in shaping the phylogeography of the Speckled Hummingbird (Adelomyia melanogenys) in Ecuador. Mol. Phylo. Evol. 43:795–807.

13. Chavez JA, Weir JT, Smith TB (2011) Diversification in Adelomyia hummingbirds follows Andean uplift. Mol. Ecol. 20: 4564–4576.

14. Barker FK, Benesh MK, Vandergon AJ, Lanyon SM (2012) Contrasting evolutionary dynamics and information content of the avian mitochondrial control region and ND2 gene. PLoS One 7:e46403.

15. Lelevier MJ (2011) Phylogeography of three widespread Neotropical avian taxa: Rufous-tailed Hummingbird, White-breasted Wood-Wren, and Anthracothorax mangos. M.S. Thesis. University of Alaska.

16. Wenner TJ, Russello MA, Wright TF (2012) Cryptic species in a Neotropical parrot: Genetic variation within the Amazona farinosa complex and its conservation implications. Cons. Genet. 13:1427–1432.

17. Eberhard JR, Bermingham E (2004) Phylogeny and biogeography of the Amazona ochrocephala (Aves: Psittacidae) complex. Auk 121:318–332.

18. Ribas CC, Tavares ES, Yoshihara C, Miyaki CY (2007) Phylogeny and biogeography of Yellow-headed and Blue-fronted parrots (Amazona ochrocephala and Amazona aestiva) with special reference to the South American taxa. Ibis 149:564–574.

19. Cuervo AM (2013) Evolutionary assembly of the Neotropical montane avifauna. Ph.D. Dissertation. Louisiana State University.

20. McCracken KG, Johnson WP, Sheldon FH (2001) Molecular population genetics, phylogeography, and conservation biology of the Mottled Duck (Anas fulvigula). Cons. Genet. 2:87–102.

21. Peters JL, Omland KE (2007) Population structure and mitochondrial polyphyly in North American Gadwalls (Anas strepera). Auk 124:444–462.

22. McCormack JE, Heled J, Delaney KS, Peterson AT, Knowles LL (2011) Calibrating divergence times on species trees versus gene trees: Implications for speciation history of Aphelocoma jays. Evolution 65:184–202.

23. McCormack JE, Peterson AT, Bonaccorso E, Smith TB (2008) Speciation in the highlands of Mexico: Genetic and phenotypic divergence in the Mexican jay (Aphelocoma ultramarina). Mol. Ecol. 17:2505–2521.

24. Ribas CC, Miyaki CY (2004) Molecular systematics in Aratinga parakeets: Species limits and historical biogeography in the ‘solstitialis’ group and the systematic position of *Nandayus nenday*. Mol. Phylo. Evol. 30:663–675.

25. Navarro-Sigüenza AG, Peterson AT, Nyari A, García-Deras GM, García-Moreno J (2008) Phylogeography of the Buarremon brush-finch complex (Aves, Emberizidae) in Mesoamerica. Mol. Phylo. Evol. 47:21–35.

26. Cadena CD, Cuervo AM (2010) Molecules, ecology, morphology, and songs in concert: how many species is Arremon torquatus (Aves: Emberizidae)? Biol. J. Linn. Soc. 99:152–176.

27. Cicero C, Koo MS (2012) The role of niche divergence and phenotypic adaptation in promoting lineage diversification in the Sage Sparrow (Artemisiospiza belli, Aves: Emberizidae). Biol. J. Linn. Soc. 107:332–354.

28. Smith BT, et al. (2014) The drivers of tropical speciation. Nature 515:406–409.

29. Puebla-Olivares F, et al. (2008) Speciation in the Emerald Toucanet (Aulacorhynchus prasinus) complex. Auk 125:39–50.

30. Cicero C (2004) Barriers to sympatry between avian sibling species (Paridae:Baeolophus) in local secondary contact. Evolution 58:1573–1587.

31. Barber BR, Klicka J (2010) Two pulses of diversification across the Isthmus of Tehuantepec in a montane Mexican bird fauna. Proc. R. Soc. B 282:2675–2681.

32. Vilaça ST, Santos FR (2010) Biogeographic history of the species complex Basileuterus culicivorus (Aves, Parulidae) in the Neotropics. Mol. Phylo. Evol. 57:585–597.

33. Gutiérrez-Pinto N,et al. (2012) Non-monophyly and deep genetic differentiation across low-elevation barriers in a Neotropical montane bird (Basileuterus tristriatus; Aves: Parulidae). Mol. Phylo. Evol. 64:156–165.

34. Pruett CL, Winker K (2005) Biological impacts of climatic change on a Beringian endemic: Cryptic refugia in the establishment and differentiation of the Rock Sandpiper (Calidris ptilocnemis). Climatic Change 68:219–240.

35. Zink RM, Kessen AE, Line TV, Blackwell-Rago RC (2001) Comparative phylogeography of some aridland bird species. Condor 103:1–10.

36. Vásquez-Miranda H, Navarro-Sigüenza AG, Omland KE (2009) Phylogeography of the Rufous-naped Wren (Campylorhynchus rufinucha): Speciation and hybridization in Mesoamerica. Auk 126:765–778.

37. González MA, Eberhard JR, Lovette IJ, Olson SL, Bermingham E (2003) Mitochondrial DNA phylogeography of the Bay Wren (Troglodytidae:Thryothorus nigricapillus) complex. Condor 105:228–238.

38. Seeholzer GF, Winger BM, Harvey MG, Weckstein JD (2011) A new species of barbet (Capitonidae:Capito) from the Cerros del Sira, Ucayali, Peru. Auk 129:551–559.

39. Kimura M, et al. (2002) Phylogeographical approaches to assessing demographic connectivity between breeding and overwintering regions in a Nearctic-Neotropical warbler (Wilsonia pusilla). Mol. Ecol. 11:1605–1616.

40. Barrera-Guzmán AO, Milá B, Sánchez-González LA, Navarro-Sigüenza AG (2012) Speciation in an avian complex endemic to the mountains of Middle America (Ergaticus, Aves: Parulidae). Mol. Phylo. Evol. 62:907–920.

41. Smith BT,et al. (2011) The role of historical and contemporary processes on phylogeographic structure and genetic diversity in the Northern Cardinal,Cardinalis cardinalis. BMC Evol. Biol. 11:136.

42. Manthey JD, Klicka J, Spellman GM (2011) Cryptic diversity in a widespread North American songbird: Phylogeography of the Brown Creeper (Certhia americana). Mol. Phylo. Evol. 58:502–512.

43. Burns KJ, Barhoum DN (2006) Population-level history of the Wrentit (Chamaea fasciata): Implications for comparative phylogeography in the California Floristic Province. Mol. Phylo. Evol. 38:117–129.

44. Oyler-McCance SJ, St.John J, Knopf FL, Quinn TW (2005) Population genetic analysis of Mountain Plover using mitochondrial DNA sequence data. Condor 107:353–362.

45. Funk WC, Mullins TD, Haig SM (2007) Conservation genetics of Snowy Plovers (Charadrius alexandrinus) in the Western Hemisphere: Population genetic structure and delineation of a new subspecies. Cons. Genet. 8:1287–1309.

46. García-Moreno J, Navarro-Sigüenza AG, Peterson AT, Sánchez-González LA (2004) Genetic variation coincides with geographic structure in the Common Bush-Tanager (Chlorospingus ophthalmicus) complex from Mexico. Mol. Phylo. Evol. 33:186–196.

47. Weir JT, Bermingham E, Miller MJ, Klicka J, González MA (2008) Phylogeography of a morphologically diverse Neotropical montane species, the Common Bush-Tanager (Chlorospingus ophthalmicus). Mol. Phylo. Evol. 47:650–664.

48. Bonaccorso E, Navarro-Sigüenza AG, Sánchez-González LA, Peterson AT, García-Moreno J (2008) Genetic differentiation of the Chlorospingus ophthalmicus complex in Mexico and Central America. J. Avian. Biol. 39:311–321.

49. Cadena CD, Gutiérrez-Pinto N, Dávila N, Chesser RT (2011) No population genetic structure in a widespread aquatic songbird from the Neotropics. Mol. Phylo. Evol. 58:540–545.

50. Chesser RT (2004) Systematics, evolution, and biogeography of the South American ovenbird genus Cinclodes. Auk 121:752–766.

51. Sanín S, et al. (2009) Paraphyly of Cinclodes fuscus (Aves: Passeriformes: Furnariidae): Implications for taxonomy and biogeography. Mol. Phylo. Evol. 53:547–555.

52. Omland KE, Tarr CL, Boarman WI, Marzluff JM, Fleischer RC (2000) Cryptic genetic variation and paraphyly in ravens. Proc. R. Soc. B 1461:2475–2482.

53. Bryson RW,et al. (2014) Diversification across the New World within the “blue” Cardinalids (Aves: Cardinalidae). J. Biogeogr. 41:587–599.

54. Smith BT, Amei A, Klicka J (2012) Evaluating the role of contracting and expanding rainforest in initiating cycles of speciation across the Isthmus of Panama, Proc. R. Soc. B 279:3520–3526.

55. Barrowclough GF, Groth JG, Mertz LA, Gutiérrez RJ (2004) Phylogeographic structure, gene flow, and species status in Blue Grouse (Dendragapus obscurus). Mol. Ecol. 7:1911–1922.

56. Cabanne GS,D’Horta FM, Meyer D, Silva J, Miyaki CY (2011) Evolution of Dendrocolaptes platyrostris (Aves: Furnariidae) between the South American open vegetation corridor and the Atlantic Forest. Biol. J. Linn. Soc. 103:801–820.

57. Isler ML, Cuervo AM, Bravo GA, Brumfield RT (2012) An integrative approach to species-level systematics reveals the depth of diversification in an Andean Thamnophilid, the Long-tailed Antbird. Auk 114:571–583.

58. Bates JM, Hackett SJ, Goerck JM (1999) High levels of mitochondrial DNA differentiation in two lineages of antbirds (Drymophila and Hypocnemis). Auk 116:1093–1106.

59. Paxton EH (2000) Molecular genetic structuring and demographic history of the Willow Flycatcher (Empidonax traillii). M.S. Thesis. Northern Arizona University.

60. Smith BT, Ribas CC, Whitney BM, Hernández-Baños BE, Klicka J (2013) Identifying biases at different spatial and temporal scales of diversification: A case study in the Neotropical parrotlet genus Forpus. Mol. Ecol. 22:483–494.

61. Dor R, Safran RJ, Sheldon FH, Winkler DW, Lovette IJ (2010) Phylogeny of the genus Hirundo and the Barn Swallow subspecies complex. Mol. Phylo. Evol. 56:409–418.

62. Fernandes AM, Wink M, Sardelli CH, Aleixo A (2014) Multiple speciation across the Andes and throughout Amazonia: The case of the Spot-backed Antbird species complex (Hylophylas naevius/Hylophylax naevioides). J. Biogeogr. 41:1094–1104.

63. Tobias JA, Bates JM, Hackett SJ, Seddon N (2008) Comment on “The latitudinal gradient in recent speciation and extinction rates of birds and mammals”. Science 319:901c.

64. Naka LN, Bechtoldt CL, Henriques LMP, Brumfield RT (2012) The role of physical barriers in the location of avian suture zones in the Guiana Shield, northern Amazonia. Am. Nat. 179:E115–E132.

65. Kondo B, Baker JM, Omland KE (2004) Recent speciation between the Baltimore Oriole and the Black-backed Oriole. Condor 106:674–680.

66. Cortes-Rodríguez N, Hernández-Baños BE, Navarro-Sigüenza AG, Omland KE (2008) Geographic variation and genetic structure in the Streak-backed Oriole: Low mitochondrial DNA differentiation reveals recent divergence. Condor 110:729–739.

67. Cortes-Rodríguez N, Hernández-Baños BE, Navarro-Sigüenza AG, Peterson AT, García-Moreno J (2008) Phylogeography and population genetics of the Amethyst-throated Hummingbird (Lampornis amethystinus). Mol. Phylo. Evol. 48:1–11.

68. Arbeláez-Cortés E, Nyári ÁS, Navarro-Sigüenza AG (2010) The differential effect of lowlands on the phylogeographic pattern of a Mesoamerican montane species (Lepidocolaptes affinis, Aves: Furnariidae). Mol. Phylo. Evol. 57:658–668.

69. Honey-Escandón M, Hernández-Baños BE, Navarro-Sigüenza AG, Benítez-Díaz H, Peterson AT (2008) Phylogeographic patterns of differentiation in the Acorn Woodpecker. Wilson J. Ornith. 120:478–493.

70. Miller MJ,et al. (2008) Out of Amazonia again and again: Episodic crossing of the Andes promotes diversification in a lowland forest flycatcher. Proc. R. Soc. B 275:1133–1142.

71. Arbeláez-Cortés E, Milá B, Navarro-Sigüenza AG (2014) Multilocus analysis of intraspecific differentiation in three endemic bird species from northern Neotropical dry forest. Mol. Phylo. Evol. 70:362–377.

72. Pérez-Emán JL (2005) Molecular phylogenetics and biogeography of the Neotropical redstarts (Myioborus; Aves, Parulinae). Mol. Phylo. Evol. 37:511–528.

73. >Pérez-Emán JL, Mumme RL, Jablonski PG (2010) Phylogeography and adaptive plumage evolution in Central American subspecies of the Slate-throated Redstart (*Myioborus miniatus*). Ornith. Monogr. 1:90–102.

74. Lovette IJ (2004) Molecular phylogeny and plumage signal evolution in a trans Andean and circum Amazonian avian species complex. Mol. Phylo. Evol. 32:512–523.

75. Batalha-Filho H, Cabanne GS, Miyaki CY (2012) Phylogeography of an Atlantic forest passerine reveals demographic stability through the last glacial maximum. Mol. Phylo. Evol. 65:892–902.

76. Miller MJ, Bermingham E, Klicka J, Escalante P, Winker K (2010) Neotropical birds show a humped distribution of within-population genetic diversity along a latitudinal transect. Ecology Letters 13:576–586.

77. Fernandes AM, Wink M, Aleixo A (2012) Phylogeography of the Chestnut-tailed Antbird (Myrmeciza hemimelaena) clarifies the role of rivers in Amazonian biogeography. J. Biogeogr. 39:1524–1535.

78. Raposo do Amaral F, Albers PK, Edwards SV, Miyaki CY (2013) Multilocus tests of Pleistocene refugia and ancient divergence in a pair of Atlantic Forest antbirds (Myrmeciza). Mol. Ecol. 22:3996–4013.

79. Dohms KM, Burg TM (2013) Molecular markers reveal limited population genetic structure in a North American Corvid, Clark’s Nutcracker (Nucifraga columbiana). PLoS One 8:e79621.

80. Zink RM,et al. (2005) Mitochondrial DNA variation, species limits, and rapid evolution of plumage coloration and size in the Savannah Sparrow. Condor 107:21–28.

81. Herr CA, Sykes Jr. PW, Klicka J (2011) Phylogeography of a vanishing North American songbird: the Painted Bunting (Passerina ciris). Conserv. Genet. 12:1395–1410.

82. van Els P, Cicero C, Klicka J (2012) High latitudes and high genetic diversity: Phylogeography of a widespread boreal bird, the Gray Jay (Perisoreus canadensis). Mol. Phylo. Evol. 63:456–465.

83. Kirchman JJ, Whittingham LA, Sheldon FH (2000) Relationships among Cave Swallow populations (Petrochelidon fulva) determined by comparisons of microsatellite and cytochrome b data. Mol. Phylo. Evol. 14:107–121.

84. van Els P, Spellman GM, Smith BT, Klicka J (2014) Extensive gene flow characterizes the phylogeography of a North American migrant bird: Black-headed Grosbeak (Pheucticus melanocephalus). Mol. Phylo. Evol. 78:148–159.

85. Campagna L,et al.(2011) A molecular phylogeny of the Sierra-Finches (Phrygilus, Passeriformes): Extreme polyphyly in a group of Andean specialists. Mol. Phylo. Evol. 61:521–533.

86. Zink RM, Rohwer S, Drovetski S, Blackwell-Rago RC, Farrell SL (2002) Holarctic phylogeography and species limits of Three-toed Woodpeckers. Condor 104:167–170.

87. Pulgarín-R. PC, Burg TM (2012) Genetic signals of demographic expansion in Downy Woodpecker (Picoides pubescens) after the last North American glacial maximum. PLoS One 7:e40412.

88. Klicka J, Spellman GM, Winker K, Chua V, Smith BT (2011) A phylogeographic and population genetic analysis of a widespread, sedentary North American bird: The Hairy Woodpecker (Picoides villosus). Auk 128:346–362.

89. Drovetski SV, Zink RM, Ericson PGP, Fadeev IV (2010) A multilocus study of Pine Grosbeak phylogeography supports the pattern of greater intercontinental divergence in Holarctic boreal forest birds than in birds inhabiting other high-latitude habitats. J. Biogeogr. 37:696–706.

90. Pravosudov VV, et al. (2012) Population genetic structure and its implications for adaptive variation in memory and the hippocampus on a continental scale in food-caching Black-capped Chickadees. Mol. Ecol. 21:4486–4497.

91. Spellman GM, Riddle B, Klicka J (2007) Phylogeography of the Mountain Chickadee (Poecile gambeli): Diversification, introgression, and expansion in response to Quaternary climate change. Mol. Ecol. 16:1055–1068.

92. Zink R, Groth JG, Vásquez-Miranda H, Barrowclough GF (2013) Phylogeography of the California Gnatcatcher (Polioptila californica) using multilocus DNA sequences and ecological niche modeling: Implications for conservation. Auk 130:449–458.

93. Valderrama E, Pérez-Emán JL, Brumfield RT, Cuervo AM, Cadena CD (2014) The influence of the complex topography and dynamic history of the montane Neotropics on the evolutionary differentiation of a cloud forest bird (*Premnoplex brunnescens*, Furnariidae). J. Biogeogr. 41:1533–1546.

94. Maldonado-Coelho M, Blake JG, Silveira LF, Batalha-Filho H, Ricklefs RE (2013) Rivers, refuges, and population divergence of fire-eye antbirds (Pyriglena) in the Amazon Basin. J. Evol. Biol. 26:1090–1107.

95. DaCosta JM, Wehtje W, Klicka J (2008) Historic genetic structuring and paraphyly within the Great-tailed Grackle. Condor 110:170–177.

96. Maley JM, Brumfield RT (2013) Mitochondrial and next-generation sequence data used to infer phylogenetic relationships and species limits in the Clapper/King rail complex. Condor 115:316–329.

97. Chaves JA, Hidalgo JR, Klicka J (2013) Biogeography and evolutionary history of the Neotropical genus Saltator (Aves: Thraupini). J. Biogeogr. 40:2180–2190.

98. Cabanne GS, Sari EHR, Meyer D, Santos FR, Miyaki CY (2013) Matrilineal evidence for demographic range expansion, low diversity, and lack of phylogeographic structure in the Atlantic forest endemic Greenish Schiffornis Schiffornis virescens (Aves: Tityridaae). J. Ornith. 154:371–384.

99. d’Horta FM, Cuervo AM, Ribas CC, Brumfield RT, Miyaki CY (2013) Phylogeny and comparative phylogeography of Sclerurus (Aves: Furnariidae) reveal constant and cryptic diversification in an old radiation of rain forest understory specialists. J. Biogeogr. 40:37–49.

100. Malpica A, Ornelas JF (2014) Postglacial northward expansion and genetic differentiation between migratory and sedentary populations of the Broad-tailed Hummingbird (Selasphorus platycercus). Mol. Ecol. 23:435–452.

101. McKay BD (2009) Evolutionary history suggests rapid differentiation in the Yellow-throated Warbler (Dendroica dominica). J. Avian Biol. 40:181–190.

102. Milot EH, Gibbs HL, Hobson KA (2000) Phylogeography and genetic structure of northern populations of the Yellow Warbler (Dendroica petechia). Mol. Ecol. 9:667–681.

103. Colbeck GJ, Gibbs HL, Marra PP, Hobson KA, Webster MS (2008) Phylogeography of a widespread North American migratory songbird (Setophaga ruticilla). J. Hered. 99:453–463.

104. Ralston J, Kirchman JJ (2012) Continent-scale genetic structure in a boreal forest migrant, the Blackpoll Warbler (Setophaga striata). Auk 129:467–478.

105. Spellman GM, Klicka J (2007) Phylogeography of the White-breasted Nuthatch (Sitta carolinensis): Diversification in North American pine and oak woodlands. Mol. Ecol. 16:1729–1740.

106. Milá B, Smith TB, Wayne RK (2006) Postglacial population expansion drives the evolution of long-distance migration in a songbird. Evolution 60:2403–2409.

107. Barrowclough GF, Groth JG, Odom KJ, Lai JE (2011) Phylogeography of Spotted Owl (Strix occidentalis): Species limits, multiple refugia, and range expansion. Auk 28:696–706.

108. Barker FK, Vandergon AJ, Lanyon SM (2008) Assessment of species limits among yellow-breasted meadowlarks (Sturnella spp.) using mitochondrial and sex-linked markers. Auk 4:869–879.

109. Stenzler LM,et al. (2009) Subtle edge-of-range genetic structuring in transcontinentally distributed North American Tree Swallows. Condor 111:470–478.

110. Rojas-Soto OR, Espinosa de los Monteros A, Zink RM (2007) Phylogeography and patterns of differentiation in the Curve-billed Thrasher. Condor 109:456–463.

111. Sgariglia EA, Burns KJ (2003) Phylogeography of the California Thrasher (Toxostoma redivivum) based on nested-clade analysis of mitochondrial-DNA variation. Auk 120:346–361.

112. Drovetski SV, et al. (2004) Complex biogeographic history of a Holarctic passerine. Proc. R. Soc. B 271:545–551.

113. Zink RM, Jones AW, Farquhar CC, Westberg MC, Gonzalez Rojas JI (2010) Comparison of molecular markers in the endangered Black-capped Vireo (Vireo atricapilla) and their interpretation in conservation. Auk 127:797–806.

114. Capurucho JMG, et al. (2013) Combining phylogeography and landscape genetics of Xenopipo atronitens (Aves: Pipridae), a white sand campina specialist, to understand Pleistocene landscape evolution in Amazonia. Biol. J. Linn. Soc. 110:60–76.

115. Aleixo A (2004) Historical diversification of a terra-firme forest bird superspecies: A phylogeographic perspective on the role of different hypotheses of Amazonian diversification. Evolution 58:1303–1317.

116. Cabanne GS, Santos FR, Miyaki CY (2007) Phylogeography of Xiphorhynchus fuscus (Passeriformes, Dendrocolaptidae): Vicariance and recent demographic expansion in southern Atlantic forest. Biol. J. Linn. Soc. 91:73–84.

117. Sousa-Neves T, Aleixo A, Sequeira F (2013) Cryptic patterns of diversification of a widespread Amazonian woodcreeper species complex (Aves: Dendrocolaptidae) inferred from multilocus phylogenetic analysis: Implications for historical biogeography and taxonomy. Mol. Phylo. Evol. 68:410–424.

118. Lougheed SC,et al. (2013) Continental phylogeography of an ecologically and morphologically diverse Neotropical songbird, Zonotrichia capensis. BMC Evol. Biol. 13:58.

